# Variant-specific interactions at the plasma membrane: Heparan sulfate’s impact on SARS-CoV-2 binding kinetics

**DOI:** 10.1101/2024.01.10.574981

**Authors:** Dario Valter Conca, Fouzia Bano, Julius von Wirén, Lauriane Scherrer, Justas Svirelis, Konrad Thorsteinsson, Andreas Dahlin, Marta Bally

## Abstract

The worldwide spread of SARS-CoV-2 has been characterised by the emergence of several variants of concern (VOCs) presenting an increasing number of mutations in the viral genome. The spike glycoprotein, responsible for engaging the viral receptor ACE2, exhibits the highest density of mutations, suggesting an ongoing evolution to optimize viral entry. However, previous studies focussed on isolated molecular interactions, neglecting the intricate composition of the plasma membrane and the interplay between viral attachment factors. Our study explores the role of avidity and of the complexity of the plasma membrane composition in modulating the virus-host binding kinetics during the early stages of viral entry for the original Wuhan strain and three VOCs: Omicron BA.1, Delta, and Alpha. We employ fluorescent liposomes decorated with spike from several VOCs as virion mimics in single-particle tracking studies on native supported lipid bilayers derived from pulmonary Calu-3 cells. Our findings reveal an increase in the affinity of the multivalent bond to the cell surface for Omicron driven by an increased association rate. We show that heparan sulfate (HS), a sulfated glycosaminoglycan commonly expressed on cells’ plasma membrane, plays a central role in modulating the interaction with the cell surface and we observe a shift in its role from screening the interaction with ACE2 in early VOCs to an important binding factor for Omicron. This is caused by a ∼10-fold increase in Omicron’s affinity to HS compared to the original Wuhan strain, as shown using atomic force microscopy-based single-molecule force spectroscopy. Our results show the importance of coreceptors, particularly HS, and membrane complexity in the modulation of the attachment in SARS-CoV-2 VOCs. We highlight a transition in the variants’ attachment strategy towards the use of HS as an initial docking site, which likely plays a role in shaping Omicron’s tropism towards infection of the upper airways, milder symptoms, and higher transmissibility.

## Introduction

Angiotensin converting enzyme 2 (ACE2) was identified as the entry receptor for severe acute respiratory syndrome coronavirus 2 (SARS-CoV-2) only weeks after the isolation of the virus^1^. Binding to ACE2 was demonstrated to be essential for successful entry, as for the closely related SARS-CoV-1 virus^2^, and its interaction with the viral glycoprotein spike has been thoroughly characterised^3,4^. However, the isolated interaction with the viral receptors is a great simplification from the reality of the first interaction between the virus and the host membrane. Indeed, the plasma membrane is a complex environment with hundreds of molecule types associated with it, from proteins to lipids and glycans^5^. A viral particle approaching the cell must navigate through plasma membrane components before binding the viral receptor^6^. The modulation of the interaction with the plasma membrane components before the receptor engagement is thus an important step that is often overseen due to the challenges arising from studying the complex and varied membrane environment^7^. Investigations of the SARS-CoV-2 entry process are further complicated by the fact that the virus has shown a great propensity to mutate, producing new variants of concern (VOCs) with increased fitness and quickly spreading through the population^8^. The highest mutation rates have been observed in the N-terminal domain and receptor-binding domain (RBD) of spike. These are the outermost and most exposed regions of spike, and the RBD is responsible for receptor engagement. This trend suggests a strong selection towards immune system evasion in a population increasingly presenting antibodies against the virus^9–12^, but also an optimisation of the interaction with the cell surface, with higher infectivity reported for emerging VOCs^13^. The emergence of the Omicron variant coincided with a particularly rapid increase in mutation in the RBD. This was also accompanied by a shift in viral tropism, with a propensity to infect the upper rather than the lower respiratory tract resulting in generally milder symptoms^14,15^. At the cellular level, a lower reliance on the TMPRSS-2 cleavage of spike and fusion at the plasma membrane in favour of endocytosis was also observed, with reports of a decreased infectivity in lung primary cultures and lung-derived cell lines (e.g. Calu-3), but not in airway cells^16^. Despite these molecular and tropism changes, the strength of the interaction between spike and ACE2 remained fairly constant among VOCs with affinities reported in the range of 0.5-50 nM^10,13,17–20^. The variation in entry tropism and the high mutation rate in the RBD, thus, cannot be simply explained by the interaction with ACE2. They instead suggest the importance of the interaction with the complex environment at the plasma membrane, including the contribution of additional attachment factors exploited by the virus whose role remains often elusive.

Several attachment factors have been proposed for SARS-CoV-2^21–27^, however, their role and interplay in the virus entry process are still to be elucidated. In particular, the interaction with heparan sulfate proteoglycans appears to be of relevance^28,29^. Heparan sulfate (HS) is a negatively charged sulfated polysaccharide with varying sulfation patterns^30^. It is commonly and widely expressed in virtually all animal tissues, where it is a major component of the cellular glycocalyx, the carbohydrate-rich meshwork of proteoglycans and glycolipids covering the cells’ plasma membrane^31^. Due to its ubiquity and charge profile, HS acts as receptor or attachment factor for numerous viruses. Virus particles usually form weak multivalent interactions that aid their movement through the glycocalyx to the cell membrane^6,7,32,33^. The interaction between SARS-CoV-2 spike and HS has been observed early in the study of the virus with Clausen et al. showing affinity to both the RBD of spike and to the full-length protein, as well as a reduction of infectivity if HS is enzymatically removed from VeroE6 cells^28^. In addition to serving as an attachment factor, HS was shown to stabilize the “open” RBD conformation needed for ACE2 engagement^28,34–36^, thereby further facilitating infection. A significant increase in the affinity to HS was observed for Omicron (BA.1), with a 3- to 5-fold increase in affinity when compared to the original Wuhan strain, possibly driven by the increased overall positive charge of spike interacting with the negatively charged HS chains^37,38^. This increase in affinity has been hypothesised to play a role in the virus tropism shift^39^. However, the study of this interaction has only been performed on isolated spike-HS pairs and without considering the complexity of the plasma membrane. In this condition, it challenging to discern the role of multivalency, avidity and the relative importance of distinct surface receptors in the overall virus interaction to the membrane. The role of HS in the attachment of SARS- CoV-2 virions to the cell surface thus remains largely unexplored at the level of a whole particle.

The direct study of the molecular interaction between SARS-CoV-2 and the plasma membrane has been hindered by the need for stable and biologically significant models for both the viral particles and the host cell surface. The use of SARS-CoV-2 viral particles requires biosafety level 3 (BSL3) conditions^40^, limiting the facilities and instruments available for study on SARS-CoV-2. Virus-like particles (VLPs)^41,42^ and pseudotyped viruses^43,44^ carrying SARS-CoV-2 spike have been developed for use in BLS2 environments. The use of such models, however, presents some drawbacks, especially for single- particle applications and for the direct comparison between VOCs. The level and uniformity of the incorporation of spike in the viral envelope is difficult to control and might differ significantly from the original virus^45^. In addition, labelling for single particle microscopy can result in high background or interference with the particle structure or binding. These problems make the characterisation of the multivalent bonds established by the whole virion particularly difficult when compared to single protein interactions. Recently, Staufer et al. proposed the use of soluble spike protein immobilised onto synthetic liposomes which resemble SARS-CoV-2 virions in size and shape^46,47^. This system, while highly simplified and not suitable for infection assays, allows for better control of the sample purity, uniformity and composition, which are important aspects in microscopy. On the membrane side, the determination of the kinetic parameters of molecular bonds requires the measurement of attachment and detachment curves in a range of concentrations of analyte^13,18^ or the observations of single particle attachment and detachment at equilibrium^48,49^. Both measurements are time-intensive and require stable conditions. Live cells are not suitable since internal cellular processes and their response to the stimulation may interfere with the membrane composition and characteristics, thus affecting binding. Fixing cells prior to measurement results in a stable substrate^50^. However, fixation can cause artificial biomolecule clustering^51^ and prevents the mobility of membrane components, making biologically relevant clustering of receptors during the interaction with the virion impossible. An alternative is reconstituting the cellular plasma membrane on a glass surface where microscopy observation with single particle resolution can be performed via total internal reflection fluorescent microscopy (TIRFM)^49,52,53^. This has been achieved using membrane blebs. When in contact with a clean glass surface, bleb rupture can be induced with the addition of synthetic lipid vesicles, producing a supported lipid bilayer incorporating the membrane components of the original cells^54,55^. In this system, protein mobility can be preserved using a polymeric spacer^56^, but the bleb composition may differ from the original membrane surface due to the nature of the blebbing process, and the protocol employed to induce blebbing^55^. Alternatively, native supported lipid bilayers (nSLBs) can be obtained from purified plasma membrane collected from mechanically disrupted cells.^57^ This method, while still needing dilution with synthetic PEGylated lipids for the formation of supported lipid bilayers, preserves the native membrane composition as well as the mobility of membrane components^58^.

In this work, we present a multiscale biophysical study elucidating the effects of avidity on the interaction between SARS-CoV-2 VOCs and the host plasma membrane, the role of HS in modulating the interaction, and how it changes across variants. We deployed a minimal-component model of the SARS-CoV-2 virion using liposomes decorated with soluble spike and employed nSLBs and TIRFM-based single-particle detection to determine the kinetic properties of the multivalent bond between VOCs and the complete plasma membrane of physiologically relevant lung epithelial cells (Calu-3). We then demonstrate the importance of HS in the regulation of the interaction with the cell membrane and report the variation in the single and multivalent affinity of spike of different VOCs to HS using immobilised HS on synthetic bilayers and atomic force microscopy-based single-molecule force spectroscopy (AFM-based SMSF).

## Results and discussion

### Spike-decorated liposomes are suitable virion mimics

We implemented fluorescent spike-decorated liposomes as virion mimics to study the role of multivalency in spike attachment to the cell membrane in a BSL1 environment. Soluble spike trimers presenting a 10xHis-tag at the C-terminus were immobilised on POPC small unilamellar vesicles (SUVs) containing rhodamine-conjugated lipids and DGS-NTA lipids, via multiple Ni-NTA/His-tag bonds, as schematically shown in Fig. 1A. The average particle diameter was measured by dynamic light scattering to be ∼125 nm, increasing to ∼145 nm after spike immobilisation, Fig. S1. This is comparable to the size of the SARS-CoV-2 envelope which measures ∼100 nm in diameter^59,60^. This procedure resulted in the production of uniform and bright particles, that can be precisely detected and tracked using TIRFM-based assays. In addition, the simplified synthetic system guarantees excellent sample purity. The stability of spike immobilisation was assessed by observing the binding and release of soluble spike onto a bilayer formed from SUVs containing a 1% molar concentration of DGS-NTA using quartz crystal microbalance with dissipation monitoring (QCM-D), as shown in Fig. 1B. Once the bond was established, extensive rinsing with PBS did not cause any significant release of spike even after more than 1 hour under constant flow. The specificity of the bond was confirmed by the near complete release of spike using imidazole, and Ni-NTA binding competitor. With the His-tag located at the C-terminus of the protein in place of the deleted transmembrane domain, the release due to imidazole competition indirectly confirmed that spike trimers are oriented as on SARS-CoV-2 virions. Being bound to lipids in the bilayer, the spike is also likely free to diffuse across the lipid surface allowing for clustering upon binding to the cell surface.

**Figure 1.**
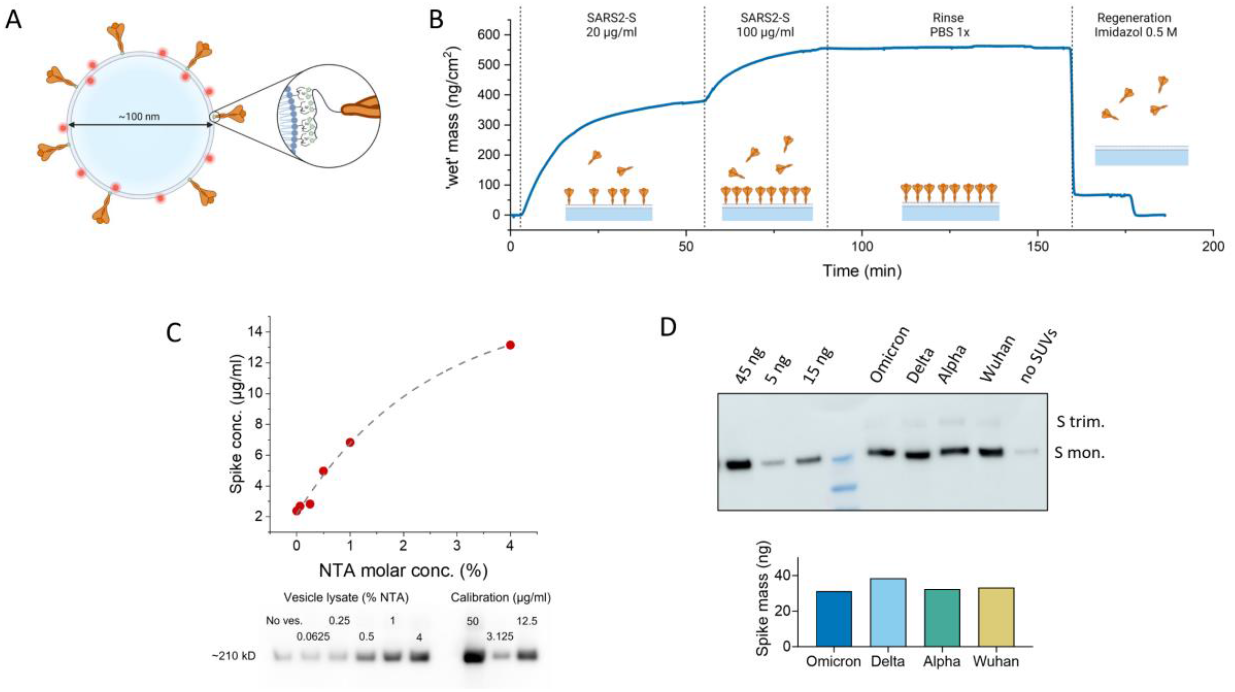
Spike-decorated liposomes are suitable SARS-CoV-2 virion mimics for single particle studies. **A**. Schematic of a spike-decorated liposome. Soluble spike, in orange, is attached to the liposomes via a multivalent bond between the NTA-conjugated lipids and the poly-His-tag located at the C-terminus of the protein. Although not shown for clarity, three 10x His-tags are present for each trimer (one per monomer). **B**. QCM-D measurement of spike attachment onto an NTA- presenting bilayer (4% molar percentage of NTA lipids). Spike trimers are injected over the sensor and incubated without flow until saturation. The formed bond is stable as no significant release is observed after rinsing in PBS for >1h. The almost complete removal of the protein by imidazole (0.5 M) indicates the specificity of the bond. The spike mass, including the water coupled with the protein layer, is estimated using the Sauerbrey model for a rigid film from the measured frequency shift. **C**. <u>Bottom</u>: Western blot immunostained for the S2 subunit of spike from the lysate of spike-decorated liposomes at molar concentrations of NTA-conjugated lipids from 0.0625% to 4% measured after Capto Core purification. Top: Plot of the spike concentration in the liposome lysate vs. the NTA molar concentration in the liposome lipid composition calculated from the western blot (red circles). The dashed grey line is the best exponential fit (*y* = *A* + *B* − *x C* where *A, B* and *C* are fitting constants) showing linearity up to 1%. **D**. Top: Western blot immunostained for the S2 subunit of spike from the lysate of NTA-liposomes for spike from Omicron, Delta, Alpha variants and the original Wuhan strain. Known amounts of spike protein (Wuhan) are loaded in the first three lanes. The last lane to the right (no SUVs) shows the soluble spike remaining in solution after filtration via Capto Core beads in the absence of NTA-liposomes. <u>Bottom</u>: Bar plot showing the quantification of the spike loading obtained from the Western blot (see methods).

We subsequently explored the possibility of controlling the spike concentration on the liposomes’ surface by adjusting the NTA content. To do so, we varied the molar concentration of NTA in the lipid mixture between 0.0625% to 4%, and quantified the spike capture efficiency of the resulting spike-decorated particles through western blot analysis, Fig. 1C. Comparing the intensity of the bands at around 210 kDa, the apparent size of glycosylated monomeric spike^61^, we observed that the spike concentration in the sample linearly depends on the percentage of NTA in the SUVs, only plateauing at concentrations higher than 1%. This was also confirmed by observing the binding of soluble spike to bilayers presenting NTA-lipids in a similar range (0.05% and 5%) using QCM-D, as shown in Fig. S2A. In addition, using surface plasmon resonance we determined that the spike density resembles that observed on SARS-CoV-2 virions (∼50 proteins/particle, see material and methods and Fig. S2B)^62,63^.

Finally, we demonstrated that the protein loading onto the liposomes is independent of the spike variant if the same tag is used. Commercial soluble spike from Omicron (BA.1), Delta (B.1.617.2), Alpha (B.1.1.7), and Wuhan strains presenting the same 10x His-tag were immobilised onto NTA-liposomes and the concentration was measured by targeting the poly-His-tag in a western blot of the particle lysate, Fig. 1D. The intensity of the resulting band varied by less than 15% between the variants, demonstrating comparable loading efficiency. Additionally, the same binding kinetics and saturation level to an NTA- presenting bilayer was confirmed for spike from Omicron and Delta variants using QCM-D, Fig. S2C.

While being an extremely simplified model of a virus particle, spike-decorated liposomes are an attractive virus mimic for single-particle studies. They allow complete control over the particle composition, both in the lipid envelope and the protein immobilised on the liposome. In this study, we employ POPC, and inert lipid, as the sole background component. While the lipid composition of the virion is more complex^46^, we opted for a simple composition to minimise interactions that are not mediated by spike, thus emphasising the differences present between spike proteins from different VOCs. Moreover, this platform serves as an exceptional tool for the systematic exploration of multivalent molecular interactions. It facilitates a direct comparison of the behaviour of various viral glycoproteins, such as spike proteins from different variants. Additionally, by using immobilised soluble spike, it allows the assessment of the multivalent binding of particles in contrast to the behaviour of isolated proteins, thus addressing the effects of avidity and clustering.

### Increased binding to pulmonary cells for Omicron

We tested the ability of spike-decorated liposomes to specifically bind to Calu-3 pulmonary epithelial cells. While many human and mammalian cell lines are susceptible to SARS-CoV-2 infection^64–66^, Calu- 3 cells were isolated from the lower respiratory tract, one of the main targets of the virus, and display an entry and replication tropism more closely related to that observed in air-water interface cultures and in vivo^16^. Cells were incubated on ice for 1 hour with spike-decorated liposomes (1% molar ratio of NTA-conjugated lipids), then washed and fixed to prevent internalisation or particle release. The cells were then imaged in epifluorescence (Fig. 2A), the bound particles were counted, and the result was normalised by the particle concentration in each sample, measured by quantification of the particles transiently entering the TIRFM excitation volume on a pure POPC bilayer, see materials and methods and Fig. S11. We observed a significant ∼2- to 3-fold increase in binding for Omicron compared to all the previous variants, as shown in Fig. 2B. No significant differences were observed between Alpha, Delta and the original Wuhan strain. In addition, particles not functionalised with spike showed negligible binding to the cell surface, excluding the presence of electrostatic effects due to the Ni-NTA positive charge. The marked increase in attachment of Omicron appears as an adaptation of the virus towards a more efficient attachment to the host plasma membrane. However, such an increase is surprising due to the reported reduced infectivity in Calu-3 cells in culture^14–16,67^. The lack of correlation between binding and infectivity suggests an increased interaction with membrane components other than ACE2. These interactions may trap the viral particle preventing efficient virus internalisation or reducing the mobility of the virus on the cell surface, thus lowering the possibility to engage ACE2. Although the contribution of later stages of viral entry, e.g., entry pathway and fusion efficiency, cannot be excluded, our observations indicate that variation in the initial binding to the cell membrane likely plays an important role in the virus tropism.

**Figure 2.**
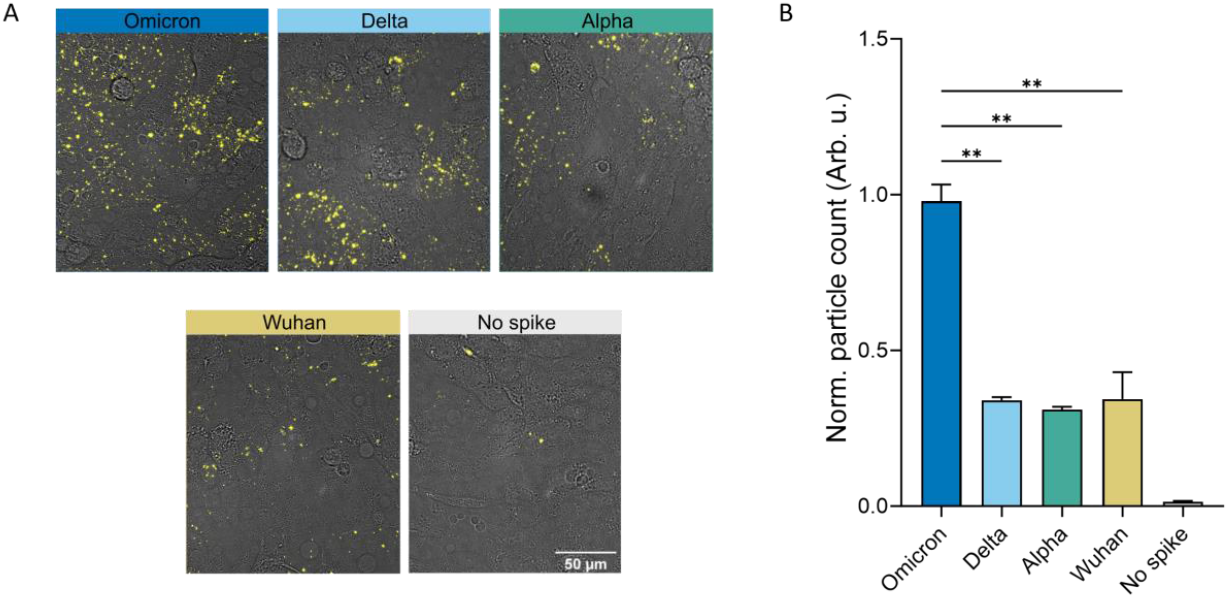
Omicron efficiently binds to Calu-3 cells. **A**. Representative brightfield and epifluorescence microscopy images of spike-decorated liposomes, in yellow, bound to Calu-3 cells, for all variants used in the study, and in the absence of spike. **B**. Relative quantification of the number of particles bound adjusted by the vesicle concentration. Data was calculated from the average of 5 replicates from two independent experiments. Statistical significance was calculated using the Brown-Forsythe and Welch ANOVA test. **: p<0.01. All variants show a p<0.001 significant increased binding with respect to SUV without spike (No spike).

### Faster association drives improved avidity for Omicron

Building on the qualitative indications of the binding assay on cells, we further characterised the dynamics of the interacting particles by investigating the binding kinetics of spike liposomes onto a native supported lipid bilayer derived from Calu-3 cells. The bilayer was formed by fusing PEGylated POPC (PEG-POPC) vesicles and native membrane material to a surface ratio of 12.5%, the highest membrane content that consistently resulted in the formation of a defect-free bilayer (data not shown). After bilayer formation, we incubated spike-decorated liposomes (1% NTA molar ratio) for at least 1 hour at 37°C to allow the system to reach equilibrium. We then performed TIRF-based equilibrium fluctuation analysis (EFA) by observing single spike-liposomes attachment and detachment from the surface^68^, as schematically shown in Fig. 3A. Detection and tracking of individual particles from the time-lapses allowed us to extract the kinetic parameters of the multivalent interaction between the spike-decorated vesicles and the nSLB. In particular, we determined the dissociation rate constant (*k*_off_), and measured the attachment rate, as shown in Fig. 3B/C. The latter, once adjusted for the particle concentration in solution, yields the measured association constant, *k*_m_, which in turn, is proportional to the association rate constant (*k*_on_). Dividing *k*_off_ by *k*_m_ yields a relative dissociation constant 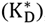, i.e., the relative difference in multivalent affinity of the system between variants. The results for all variants considered in this study are shown in Fig. 3D-F normalised by the value measured for Omicron. We report a negligible attachment in the absence of spike, confirming that the system has low non-specific signal. Most interestingly, we observed a significant increase in the association rate of Omicron, compared to all the other variants tested, except for Alpha. No significant difference between variants was observed for *k*_off_, although Omicron displayed the slowest dissociation rate on average and an increased fraction of particles that remain irreversibly linked to the substrate during the length of the experiment (irreversible fraction), Fig. S3A. Similar dissociation rates may be due to the high-affinity interaction with the ACE2 receptor for all SARS-CoV-2 variants^13,18,28^. Indeed, we speculate that after the initial attachment, possibly through several attachment factors, the particle creates a stable bond with ACE2, which dominates the detachment kinetics. In addition, the creation of multiple bonds to the membrane, which stabilise the interaction and strongly reduces detachment (the irreversible fraction is ∼50% for all variants, Table S1), might mask differences between variants in the detachment kinetics with possible coreceptors. Combining association and dissociation rates, we observed a ∼3-fold increase in affinity of the multivalent interaction for Omicron when compared to Wuhan and Delta, with a trend indicating an advantage over Alpha as well, although without statistical significance (Fig. 3F). The magnitude of the affinity increase is comparable to the increased binding observed on Calu-3 cells for Delta and Wuhan. The high multivalent affinity to the nSLB shown by Alpha, similar to what was observed for Omicron, may be the result of a higher affinity of Alpha’s RBD to ACE2 than that of the other VOCs, as reported by Han et al.^13^. The incorporation of synthetic lipids in the nSLB, indeed, dilutes the membrane material, possibly increasing the accessibility of ACE2 compared to the interaction with the cell surface. All in all, our results show an increased affinity for the multivalent interaction between Omicron to the plasma membrane of lung epithelial cells and suggest an important and evolving role of co-receptors and ACE2 accessibility.

**Figure 3.**
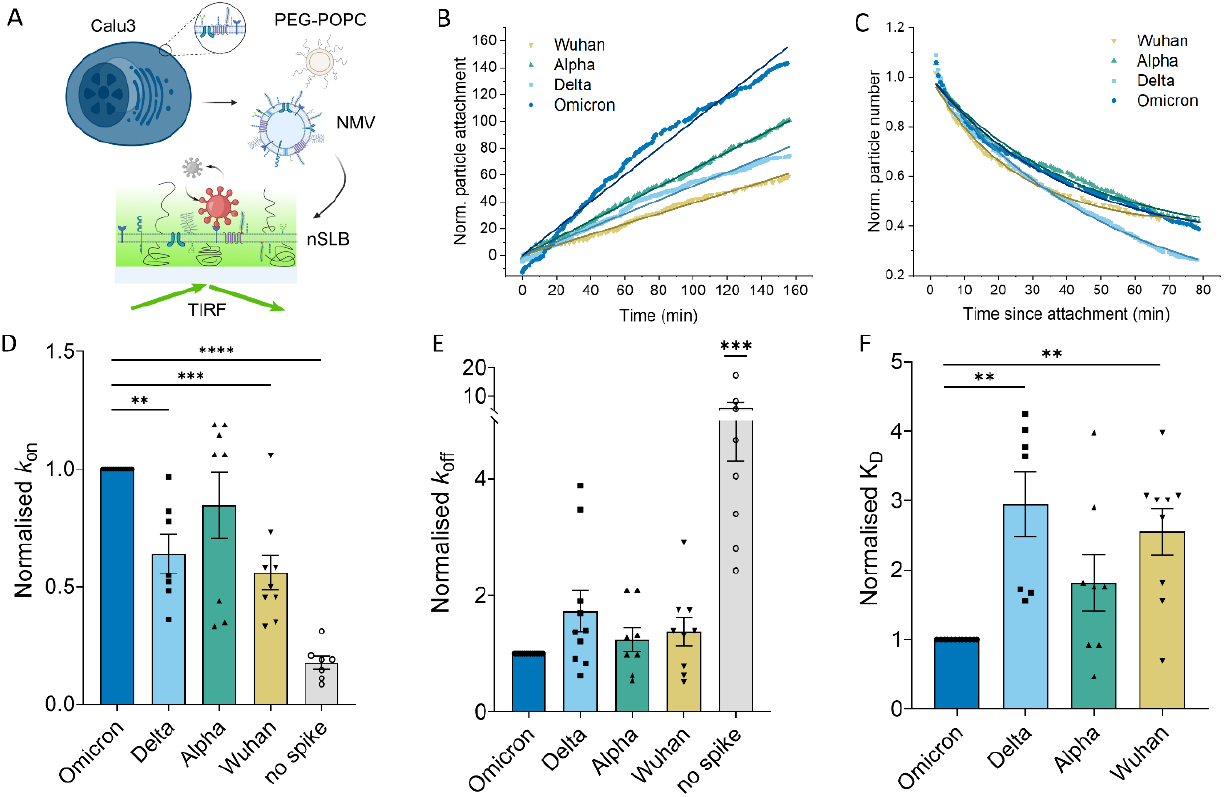
Omicron improved multivalent affinity to plasma membrane extracts is driven by increased attachment. **A**. Sketch showing the formation of a native supported lipid bilayer (nSLB). Membrane material is extracted from Calu-3 cells and purified to form native membrane vesicles (NMVs). NMVs are then mixed via sonication with PEG-POPC via sonication and then allowed to rupture on clean glass to form nSLBs. The attachment of spike-decorated vesicles on the nSLB is imaged using TIRF microscopy. **B**. Example graph of the cumulative attachment of spike-decorated vesicles from different variants with the linear fit used to extract the association rate. **C**. Example graph of the number of particles attached to the surface vs. the time elapsed since their attachment. The solid line indicates the single exponential fit with offset used to calculate *k*_off_. **D-F**. Kinetic parameters (*k*_on_, *k*_off_ and K_D_ in D, E and F respectively) measured via equilibrium fluctuation analysis of the interaction between spike-decorated vesicles and Calu-3-derived nSLBs. Data are normalised by the value measured for Omicron in each experiment. After normalisation, *k*_on_ and K_D_ are equivalent to the measured constants *k*_m_ and 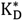 . A summary of data before normalisation is presented in Table S1. Statistical significance was calculated using one-way ANOVA test. **: p<0.01, ***: p<0.001, ****: p<0.0001. In E, *** indicates the lowest significance between the negative control and all the other conditions tested.

### Heparinase treatment increases binding to plasma membrane extract for all VOCs but Omicron

We aimed to identify the main factor responsible for the increase in interaction with the plasma membrane and proceeded with enzymatically removing HS from the nSLB surface using a mix of Heparinase I and III, Fig. 4A. HS has been shown to interact with spike and it is believed to be an important attachment factor for SARS-CoV-2^28,69–71^. Successful HS removal from the bilayer was verified by flow cytometry of antibody-stained nSLBs formed on glass beads, Fig. S4. The result of the kinetic assay on heparinase-treated bilayers is shown in Fig. 4B-D. As in the untreated case, we observed the main differences in the arrival rate with a significantly higher *k*_m_ for the Alpha variant compared to all the other tested variants but Wuhan. Omicron, which had the highest attachment rate on untreated nSLBs, now showed a lower association than Wuhan and Alpha (2- and 3-fold reduction, respectively) and a similar level to Delta, Fig. 4B. This was also reflected in a trend towards higher 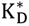, i.e., lower multivalent affinity, than Alpha, Fig. 4D. As for the untreated nSLBs, no clear differences were observed in particle dissociation, except for particles lacking spike, Fig. 4C. In this case, similar irreversible fractions were also measured for all variants, Fig. S3B. By comparing the relative association rate with or without the enzymatic treatment of the nSLB, Fig. 4E, we show that Omicron binding was not significantly affected by HS removal. Conversely, the attachment rate was increased by the enzymatic treatment for early variants. This resulted in a >2-fold increase in affinity of the multivalent interaction for Alpha and Wuhan as shown in Fig. 4F. The Delta variant also displayed a similar trend, with a ∼40% increase in the association rate constant after heparinase treatment, albeit not statistically significant. These results suggest that HS chains on the cell surface hide high-affinity receptors from the virus and partially prevent the interaction with ACE2, or other possible attachment factors on the cell surface. The direct interaction observed in the literature between spike and HS is not sufficient to compensate for the masking of ACE2^28,34^. The increased accessibility of ACE2 after the enzymatic removal of HS is also supported by the high affinity measured for Alpha, which has been reported to have the highest affinity to ACE2 of the VOCs considered in the study^13^. Previous studies showed a reduction in infection upon HS removal for SARS-CoV-2 spike pseudotyped viruses^28^ and early SPR measurement demonstrated interaction between Wuhan spike and HS^28^. Our results portray a more complex role of HS in SARS-CoV-2 infection than previously reported, with what could be a regulatory effect of the virus-host interaction. We speculate that HS acts as a weak but abundant initial attachment factor, but also reduces the frequency of binding to ACE2. This is less the case for Omicron, for which the surface interaction seems not to be altered by the presence of HS. This behaviour is most likely caused by a stronger interaction with HS which compensates for the masking effect observed in the other variants.

**Figure 4.**
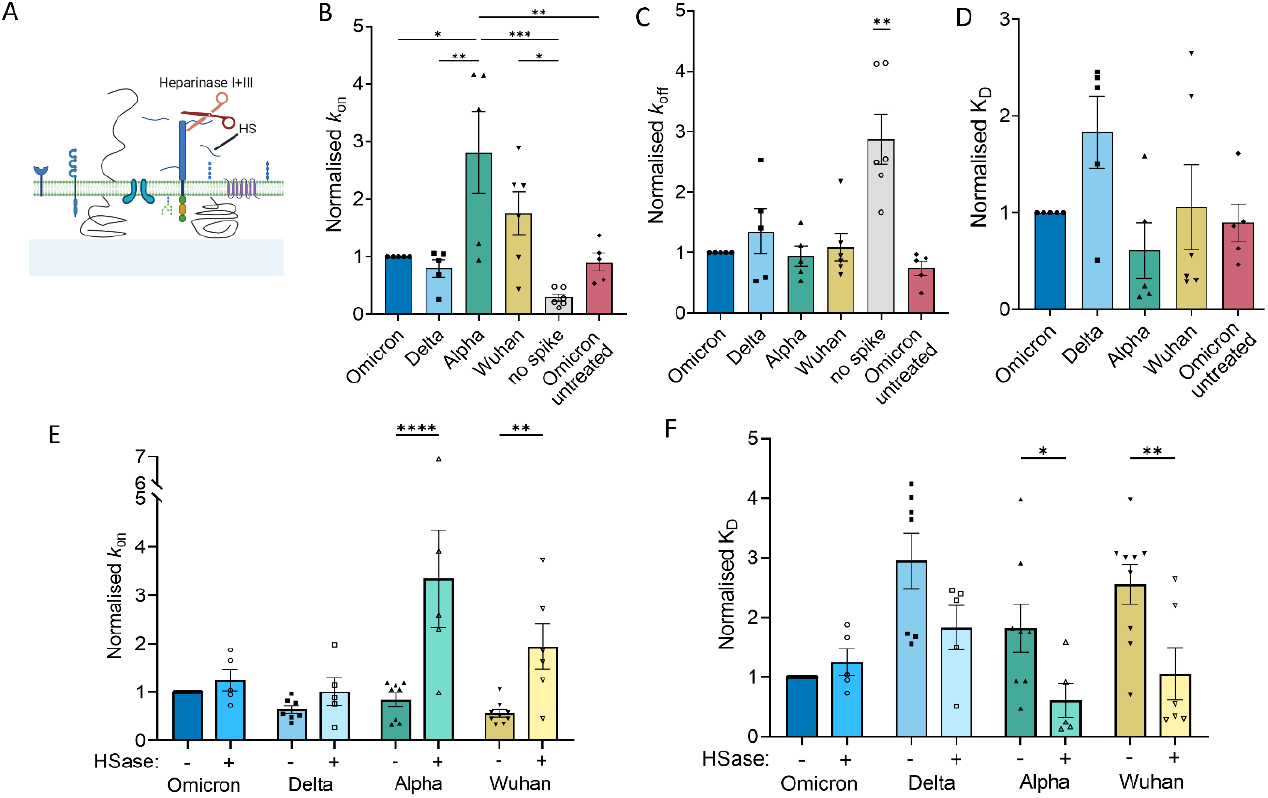
Removal of HS improves binding to nSLBs for all variants but Omicron. **A**. Schematic representation of the heparinase I+III treatment on a formed nSLB. **B-D**. Kinetic parameters (*k*_on_ in B, *k*_off_ in C and K_D_ in D) measured for each variant after the treatment with the heparinase I+III cocktail and Omicron on an nSLB mock-treated with PBS. Values are normalised in each experiment to Omicron. A summary of data before normalisation is presented in Table S2. **E-F**. Association rate constant (E) and dissociation constant (F) for each variant, with or without enzymatic removal of HS (HSase). Statistical significance is calculated only within each variant. Statistical significance was calculated using one-way ANOVA test. *: p<0.05, **: p<0.01, ***: p<0.001. In C, ** indicates the lowest significance between the difference in the negative control and all the other conditions tested.

### Increased multivalent affinity to HS for more recent variants

Noting the effect of HS on the binding kinetics on cell membrane extracts, and the previous reports of interaction between HS and purified spike from VOCs, we tested the multivalent interaction of spike-decorated liposomes to surface-immobilised HS. We formed a synthetic biotinylated supported lipid bilayer and conjugated unfractionated HS extracted from porcine mucosa through a streptavidin bridge, Fig. 5A. Hyaluronic acid, a non-sulfated but negatively charged GAG, was used as a negative control. The strong reduction of binding observed on HA indicates that the interaction to HS is not purely driven by the electrostatic attraction between the positively charged spike to the negatively charged substrate, and that specificity to the HS structure and sulfation pattern is likely an important determinant of the interaction. EFA showed an overall tendency towards a faster attachment to HS of more recent variants. Omicron, and to a lesser extent Delta, displayed a significantly faster attachment with an increment in *k*_m_ of ∼10 times for Omicron and ∼3 times for Delta when comparted to earlier variants, Fig. 5B. The increase in association rate is accompanied by an increase in the equilibrium surface coverage, indicating an overall higher multivalent affinity to HS for Omicron and Delta, Fig. 5C. Here, we could not obtain a value for the dissociation constant, as the interaction between the particles and the surface proved to be too stable, resulting in too few detachment events to confidently measure the dissociation rate constant over the duration of the experiment. This result confirms the increase in interaction between spike and HS in newer variants and it is in agreement with previous reports between isolated Omicron spike and HS^37,38,72^. The differences are, in our case, likely enhanced by the multivalency of the bond due to the presence of multiple spikes on the same particle.

**Figure 5.**
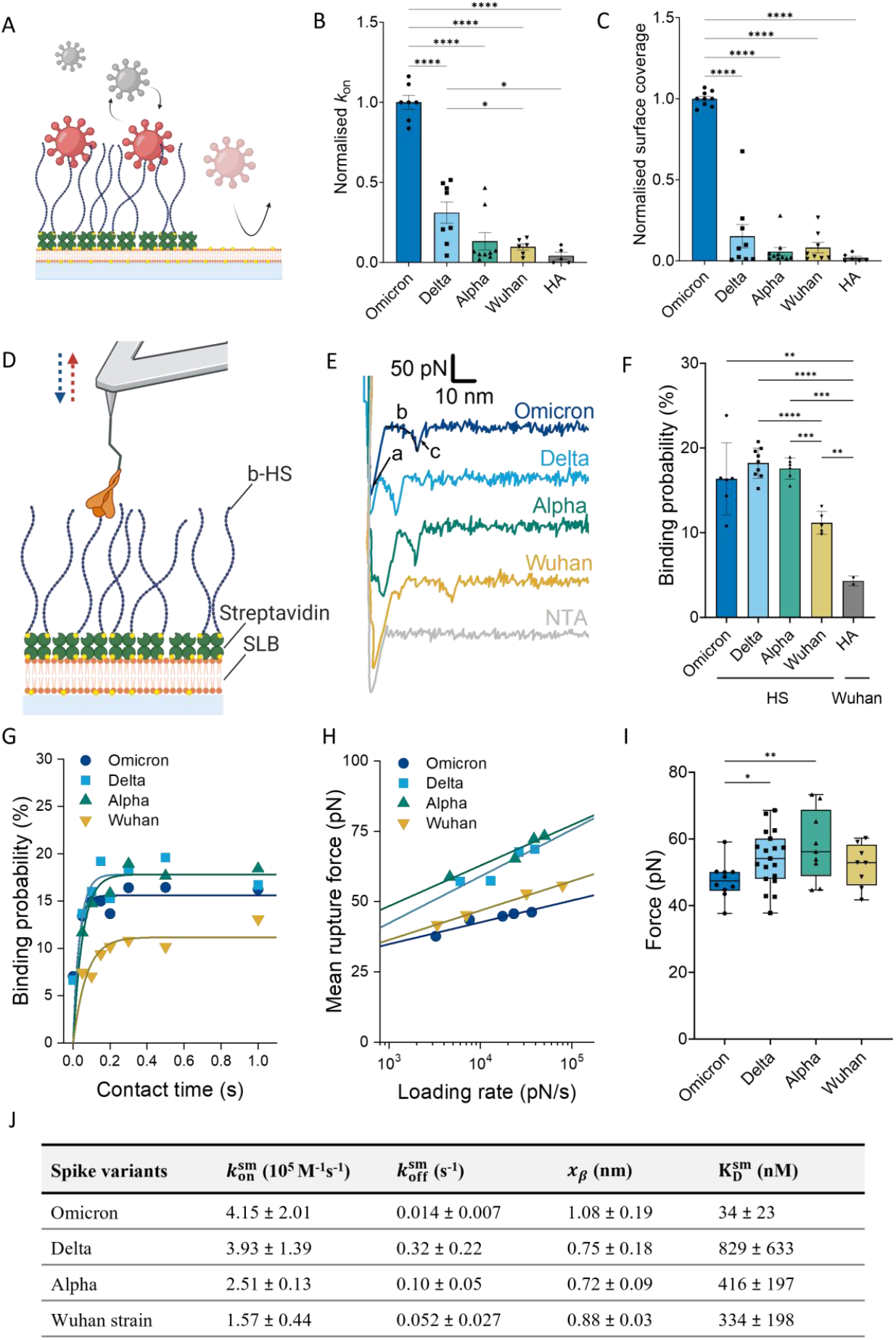
Large variations in HS binding kinetics between variants at the single-particle and single-molecule level. **A**. Schematic of the immobilization of biotinylated HS to a biotin-presenting supported lipid bilayer via a streptavidin bridge. This allows the direct observation of the binding of spike-decorated vesicles to HS via TIRF microscopy. **B**. Association rate constant for vesicles functionalised with spike from different variants. **C**. Average number of particles bound to the HS surface at equilibrium. A summary of data before normalisation is presented in Table S3. **D**. Schematic representation of AFM-SMFS setup (not to scale) to quantify the binding interaction between SARS-COV-2 spike variants and HS. **E**. Exemplary force-distance curves to display specific unbinding events. (a) Unspecific peak, (b) PEG stretching, and (c) spike-HS bond rupture. **F**. Binding probability in percentage fraction at retract velocity of 1 µm/s and surface dwell time of 0.3, 0.5 and 1s from at least 2 independent experiments to identify the specificity of spike variants binding to HS. **G**. Representative data on the influence of surface dwell time on the binding probability of spike variants and HS interactions at a fixed retract velocity of 1 µm/s. Data fit with a single exponential function for pseudo-first-order binding kinetics (solid line) to extract the association rate constant, 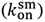, see supplementary figure S8 for the overview of all data sets. **H**. Representative dynamic force spectra (mean rupture force vs. loading rate) at a fixed dwell time of 0 s for four indicated spike variants. Fitting according to the Bell-Evans model (solid lines) yields the dissociation rate constant 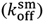 and the barrier width (*x*_β_) of a single energy barrier, see supplementary figure S9 for the overview of all data sets. **I**. Average rupture force over the loading rate range, reflecting the binding strength of individual spike-HS bonds. Each point represents the average over each loading range for every sample tested. **J**. Summary of the single molecule association 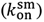 and dissociation 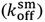 rate constants, energy barrier width (*x*_β_) and binding affinity (K_D_) of spike variants for HS measured by AFM-SMFS. P-values were determined by one-way ANOVA test: *: p<0.05, **: p<0.01, ***: p<0.001, *\*\*\*\*:* p<0.0001.

### AFM-based SMFS confirms evolving interaction to HS in variants

Given evidence of both HS binding to spike, its screening effect on nSLBs, and the significant variation in the HS role between variants, we characterised the spike-HS bond using AFM-based SMSF, Fig. 5D. In these experiments, the AFM cantilever is functionalised with a very low density of spike proteins immobilized on the tip apex through a flexible PEG linker. The tip is brought into contact with the HS surface and allowed to form bonds. Upon retraction of the cantilever at a set speed, an increasing force is applied to the bond, eventually causing it to rupture^73^. Both the distance to the surface and the force exerted by the AFM tip can be simultaneously measured, yielding force-distance (FD) curves. An exemplary FD curve for each variant is shown in Fig. 5E. Such a curve exhibits a non-specific adhesion peak followed by the characteristic stretching of PEG, and finally the bond rupture event. The rupture events were specific to spike-HS interactions as they were largely absent for the NTA-coated tip (i.e., in the absence of spike) against the HS surface (light grey FD curve in Figure 5E, light grey bar in Fig. S7) and for spike-coated tips (Wuhan) against HA (grey bar in Fig. 5F). In addition, we verified that spike-HS interactions were mainly originating from single unbinding events through several observations: (i) the binding probability (BP) was <20% for spike-HS bonds^74,75^, Fig. 5F; (ii) the Kuhn length of the PEG linker was 0.68 ± 0.13 nm, which is in agreement with the literature value of 0.7 nm for the stretching of a single PEG chain^76^, Fig. S5; and (iii) all force histograms had a unimodal Gaussian distribution, Fig. S6.

From the SMFS data, we computed the kinetic parameters of the spike-HS interaction, which are summarised in the table in Fig. 5J. The single-molecule association rate constant 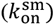 was obtained by studying the increase in binding probability with contact time, Fig. 5G and Fig. S8. We observed a monotonic increase of the attachment rate with the time of emergence between variants (Omicron>Delta>Alpha>Wuhan) in good agreement with the binding kinetics for spike-decorated liposomes to HS films. The single-molecule dissociation rate constant 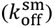, was determined by plotting the unbinding forces as a function of instantaneous loading rate (the effective linker stiffness multiplied by the retract speed, see material and methods) and fitting them to the Bell-Evan model, Figure 5H and Fig. S9. Omicron showed the lowest dissociation rate. However, in this case, a more complicated interaction was observed for the other variants, with higher 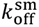 values, i.e., a faster dissociation, for Delta and Alpha compared to Wuhan (∼6 and ∼2 times respectively). Taken together, Omicron formed a considerably more stable bond with HS, as evidenced by a single-molecule dissociation constant 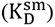 10 times lower values for Omicron with respect to reference Wuhan (Fig. 5J). Surprisingly, 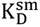 for Delta and Alpha was ∼2.5 and ∼1.2 times higher than Wuhan, respectively. These results indicate significant differences in the interaction characteristics of the various variants with HS, with a more dynamic bond for Alpha and Delta, and a stable high-affinity interaction for Omicron.

Previous studies using ELISA or SPR and soluble spike or the isolated RBD showed a small increase in the affinity of Delta to HS, similar to the results of our TIRFM assay, and no clear indication of significant variations in the attachment and detachment rates compared to the Wuhan strain^37,38^. The reason for this discrepancy might lie in the technique used. When measuring the adsorption of proteins onto a functionalised surface, the protein can interact with more than one HS chain, or establish multiple bonds with the same chain, masking possible rapid creation and rupture of single bonds. Using AFM- based SMFS, we directly addressed single bonds, not allowing possible rebinding and preventing avidity effects^77^. The simultaneous increase of attachment and detachment rates indicates that variants before Omicron did not evolve to increase the affinity to HS but instead exhibited a more dynamic interaction. We speculate that they use HS as a first attachment point, hence the higher attachment rate, but only produce transient bonds. This could allow the viral particle to more easily move through the cellular glycocalyx and transfer to the ACE2 receptor, as it has been proposed for other viruses^6^, or even move more efficiently to the lower respiratory tract and the lungs. Notably, in addition to more dynamic binding behaviour, Alpha and Delta showed a significantly higher unbinding force over the whole range of loading rate probed compared to Omicron (Fig. 5I). A high mechanical stability without correlation with the binding affinity, has been previously demonstrated for CD44 and hyaluronan interaction^75,76,78^. In our case, it might be of advantage when considering the shear forces present in the respiratory tract^79,80^, allowing the virus to stay attached to the host cell while dynamically navigating the glycocalyx to finally engage ACE2.

In the case of Omicron, the affinity is increased to ∼30 nM, similar to that of the interaction with ACE2. This results from both the fastest attachment rate and slowest dissociation measured for all the VOCs tested. This shift in the tropism towards an increased reliance on HS can be seen as one of the causes for the changes in the symptoms observed in Omicron infection and the increased transmissibility of the virus. Heparan sulfate is almost ubiquitously expressed in tissues and highly expressed both in the nasal cavity and in the lungs^71^. We speculate that the availability of binding sites in the nasal cavity and upper respiratory tract prevents the virus from travelling far in the airways and reaching the lungs in large amounts. This causes an infection of the upper respiratory tract with lower chances of lung inflammation, and an overall milder disease. This could also represent an evolutionary advantage for the virus as the infection closer to the nose and mouth results in more efficient incorporation of viral particles in aerosols^81^, causing high infectious shedding and ultimately highly transmissibility^67^.

## Conclusions

The emergence of new SARS-CoV-2 variants has been characterized by the accumulation of mutations in the spike proteins and the RBD. This has been shown to alter the interaction with ACE2 and HS, which possibly relates to changes in infectivity, transmissibility and tropism between VOCs. However, the characterisation of these interactions is often performed using single isolated receptors in simplified systems, making it difficult to quantify the relative importance of different surface receptors. In our study, we present the first characterisation of the kinetic properties of the multivalent bond formed by particles presenting SARS-CoV-2 spike proteins and the plasma membrane of host cells in the early stages of the virus entry and highlight how this interaction evolved with the emergence of new variants. We start from observations in a highly complex system, the binding of virion mimics on Calu-3 cells. We then progressively reduce the complexity of the system to allow an in-depth characterisation of the stability and kinetics of the bond formed, both at the single-particle and single-molecule level, elucidating the role of HS for several VOCs.

After validating and characterising spike-decorated liposomes as virion mimics for single-particle studies, we confirm a general tendency in the strengthening of the interaction to the plasma membrane of biologically significant cells in successive variants and the importance of the interaction of the spike protein with HS proteoglycans present on the cell surface. We elucidate the role of HS in the multivalent interaction with the cell surface, showing how, with the emergence of Omicron (B1.A), the increase in affinity to HS resulted in a marked shift in the molecule’s role. Despite spike showing affinity towards HS since the emergence of SARS-CoV-2 in Wuhan, our study highlights that upon interaction with the complex plasma membrane, the presence of HS can result in a reduction of the overall binding to the cell surface probably due to the masking of the ACE2 receptor. This is not the case for Omicron for which HS acts as a high-affinity coreceptor. In addition, AFM-based single molecule force spectroscopy measurements highlight a high-affinity interaction between Omicron’s spike and HS, contrasting with the mechanically strong but dynamic one observed in previous VOCs.

Our study improves the understanding of the role of HS in SARS-CoV-2 binding and evolution, by addressing both the molecular properties of the interaction with spike and the role that they play in virus attachment. It gives insights into how viral attachment can be linked to a shift in the tropism of VOCs and the resulting symptoms of the infection. It also shows the importance of accounting for the complexity of the plasma membrane when investigating the role of membrane components in viral attachment, as well as the synergy necessary between cellular and molecular models and techniques. In doing so, we developed biophysical techniques and remarked on the importance of multiscale and transversal studies to elucidate the biological role of physical and chemical interaction. Given the flexibility of our platform and approach, we envision our study to be expanded in the future to address other proposed coreceptors and new emerging variants, building towards a complete picture of the concurring molecular interactions exploited by SARS-CoV-2.

## Acknowledgements

The authors would like to thank the Biochemical Imaging Center at Umeå University and the National Microscopy Infrastructure, NMI (VR-RFI 2019-00217) for providing assistance in microscopy and Elliot Eriksson for the help with the calibration of the “bouncing particle analysis”. All schematic in this paper figures were created with BioRender.com. This project has been funded by the Kempe Foundation, the Knut and Alice Wallenberg Foundation, the Swedish Research Council (2017-04029; 2020-06242) and the European Union’s Horizon 2020 research and innovation programme under the Marie Skłodowska-Curie grant agreement No 101027987.

## Supplementary information

Supplementary information contains material and methods, supplementary figures referenced in the main text regarding spike-decorated liposomes and nSLB characterisation, additional kinetic data from EFA and AFM-based SMFS, and the summary of non-normalised data from EFA.

