## Supplementary methods, figures and tables for "Variant-specific interactions at the plasma membrane: Heparan sulfate’s impact on SARS-CoV-2 binding kinetics"

#### Content

Material and methods *1*

Supplementary references 12

Supplementary figures 14

Supplementary tables 19

#### Material and methods

**Small unilamellar vesicles production**

1-palmitoyl-2-oleoyl-glycero-3-phosphocholine (POPC, 850457P), 1,2-dioleoyl-sn-glycero-3-[(N-(5-amino-1-carboxypentyl) iminodiacetic acid) succinyl] (DGS-NTA, 790528P), 1,2-dioleoyl-sn-glycero-3-phosphoethanolamine-N-(lissamine rhodamine B sulfonyl) (Liss-Rhod, 810150C), 1,2-dioleoyl-sn-glycero-3-phosphoethanolamine-N-(capbiotinyl) (bioDOPE, 870273) and N-palmitoyl-sphingosine-1-{succinyl[methoxy(polyethylene glycol)5000]} (PEG-cer, 880280P) were purchased from Avanti Polar Lipids (Alabaster, AL, USA) and Oregon Green™ 488 1,2-Dihexadecanoyl-sn-Glycero-3-Phosphoethanolamine (OG-DHPE, O12650) from Thermo Fisher Scientific (Waltham, MA, USA). All lipids were received in lyophilised form and stored at -20ºC. To produce small unilamellar vesicles (SUVs), the lipids were dissolved in chloroform at concentrations between 2 and 50 mg/ml, mixed to the desired molar concentration and dried first under gentle nitrogen flow and then under vacuum overnight. They were then resuspended in 1 ml phosphate buffer saline (PBS, Medicago AB, Sweden), or HEPES buffer saline (HBS, 140 mM NaCl, 10 mM HEPES) if containing NTA groups, at a concentration between 0.5 and 2 mg/ml. SUVs were formed by extruding at least 21 times through a polycarbonate membrane, using a mini extruder (Avanti Polar Lipids; 610020), as previously described[1]. Pure POPC vesicles, POPC:bioDOPE (95:5 molar ratio), POPC:PEG (99:1 molar ratio) and POPC:OG-DHPE (99:1 molar ratio) used for the creation of supported lipid bilayers (SLB) were extruded through a 50 nm membrane. For the production of spike-decorated liposomes, POPC:DGS-NTA:Liss-Rhod (99-X:X:1 molar ratio, where X is the percentage of DGS-NTA lipids) were flash frozen and thawed 5 times by alternate immersion in liquid nitrogen and 37ºC water bath and then extruded at least 21 times through a 100 nm membrane. All stocks were stored at 4ºC for up to three months.

**Glycosaminoglycans and proteins**

Heparan sulfate purified from porcine mucosa (HS; GAG-HS01 BN1) was purchased from Iduron (UK). Lyophilized end-biotinylated HA (b-HA) was kindly provided by Innovent e.V., Technologieentwicklung (Jena, Germany). All lyophilized HS samples were dissolved by gently mixing overnight at 4°C in milli-Q (Millipore integral system, Molsheim, France) $H_2O$ to form a stock concentration of 25 mg/ml, while b-HA was prepared at the stock concentration of 2 mg/ml. The HS was biotinylated at its reducing end by oxime oxidation as described previously in [2]. The final products were aliquoted and stored at -20°C.

Lyophilized streptavidin (Sigma Aldrich, Burlington, MA, USA) was dissolved in milli-Q at 5 mg/ml, aliquoted, and stored at -80ºC. Soluble 10xHis-tagged SARS-CoV-2 spike stabilised to retain the prefusion conformation, was acquired lyophilised from ACROBiosystems (Newark, DE, USA) for all variants (Product number: original strain, Wuhan: SPN-C52H9; Alpha, B.1.1.7: SPN-C52H6; Delta, B.1.617.2: SPN-C52He; Omicron, BA.1: SPN-C52Hz), resuspended at 0.66 mg/ml in milli-Q water and 0.1% BSA, aliquoted in 4.5 µl, flash-frozen in liquid nitrogen, and stored at -80°C.

**Spike functionalisation of NTA-liposomes**

On the day of the experiment, POPC:DGS-NTA:LissRhod vesicles were diluted 1:4 (vol) in HBS. $NiCl_2$ was added to a final concentration of 20 mM for a final volume of 100 µl and the vesicles were incubated in the dark at room temperature for 30 minutes. $NiCl_2$ was removed using MicroSpin™ S-200 HR columns (27512001, Cytiva, Sweden). In brief, the columns were first washed 3 times by adding 300 µl of HSB and spinning at 700×g for 45 s. After washing, the vesicle solution was added to the column, and the elute was collected by centrifugation at 700×g for 1 min. A spike aliquot was thawed and 2 µl was added to 20 µl of the activated vesicle solution to a final concentration of 60 µg/ml. After mixing, the sample was incubated at room temperature in the dark for 2 hours. Spike aliquots were never refrozen and were kept at 4ºC for a maximum of 3 days. After incubation, the unbound spike was removed by incubation with Capto Core 700 beads (17548102, Cytiva). 100 µl of beads stock solution was washed 3 times in 1 ml PBS and collected by centrifugation at 1,000×g for 1 min. The beads were then resuspended in 50 µl of PBS and 20 µl was added to the vesicle solution. We incubated the solution for 30 min at 4°C under gentle rotation. The solution was then centrifuged at 1,000×g for 1 min to separate the beads from the vesicles, and the supernatant was run through MicroSpin™ S-400 HR columns (27514001, Cytiva) using the same protocol as for MicroSpin™ S-200 HR columns with a centrifugation speed of 600×g. The elute containing spike-decorated liposomes was collected and stored at 4ºC until use the same day.

**Characterisation** of **the size distribution** of **spike-decorated liposome**s **via DLS**

5 µl of spike-decorated liposomes were collected after incubation with Capto Core 700 beads and diluted to a final volume of 50 µl in PBS. The solution was added to a 40 µl cuvette (ZEN0040, Malvern Panalytical Ltd, UK) and measured using dynamic light scattering with a Zetasizer Nano S (Malvern Panalytical Ltd). The data were analysed using Zetasizer proprietary software (Malvern Panalytical Ltd). Two independent measurements were performed for each sample.

**Spike quantification via western blot**

The amount of spike captured by NTA-liposome was quantified via a western blot assay. In brief, 20 µl of spike-conjugated liposomes was mixed with 4 µl of SDS and 1 µl of 1 M imidazole to dissolve the vesicles and detach the spike. 5 µl of Laemmli buffer was added and the samples were incubated at 95ºC for 10 min. The samples were then carefully loaded into a precast 15-well bis-tris SDS gel (NW04125BOX, Thermo Fisher Scientific) and run at 200 V for 35 min in 5% NuPAGE™ MES SDS Running Buffer 20X (NP0002, Thermo Fisher Scientific) in milli-Q. A PVDF (polyvinylidene difluoride) membrane was prewetted in methanol and proteins were transferred in transfer buffer solution (10% 10x Tris/Glycine Buffer (1610734, Biorad), 20% Methanol and 70% milli-Q $H_2O$) using a constant current of 300 mA for 45 min in a Mini Trans-Blot® Cell (Biorad). The membrane was blocked in 5% fat-free milk in PBS + 0.05% Tween-20 (PBST) for 1 hour and then incubated with anti-SARS-CoV-2 S2 antibody (MA5-35946, Thermo Fisher Scientific) diluted 1:1000 in 5% milk in PBST at 4ºC overnight. After 3 washes in PBST, the membrane was incubated in a 1:2000 dilution of horseradish peroxidase (HRP) conjugated anti-mouse antibody (AB_228307, Invitrogen) in 5% milk in PBST. Finally, the membrane was thoroughly washed and the chemiluminescence signal was imaged using SuperSignal™ West Pico PLUS chemiluminescence substrate (Thermo Fisher Scientific) in an Amersham™ Imager 680 blot reader (Cytiva, Chicago, USA). The spike content for each variant was estimated by measuring the band intensities at a MW of ~210 kDa using an existing plugin in ImageJ. A calibration curve was obtained by the serial dilution (3x) of soluble spike (Delta) and used to convert from the band intensity to the mass of protein contained in each sample (Fig. S10).

**Multi-Parametric Surface Plasmon Resonance**

To obtain silica-coated chips for SPR, SPR chips with ~2 nm chromium (Cr) and 50 nm gold (Au) were prepared by electron-beam-heated physical vapor deposition (Lesker PVD 225) on glass substrates (20 x 12 mm; BioNavis), cleaned beforehand with RCA-2 (1:1:5 vol. conc. of HCl:$H_2O_2$(30%):Milli-Q $H_2O$) at 80°C and 50 W $O_2$ plasma at 250 mTorr. An additional thin silica layer (~11 nm) was deposited on top of the gold layer with atomic layer deposition (Oxford FlexAL) at 300°C by using a bis(t-butylamino)silane (BTBAS) as a precursor and oxygen as processing gas. Sensor surfaces were incubated in a UV/ozone chamber for at least 30 min before use.

The SPR Navi$^{TM}$ 220A instrument (BioNavis) is equipped with three lasers (wavelengths of 670 nm, 785 nm, and 980 nm), which induce different decay lengths of sensing fields. The SPR spectra can be acquired in parallel from 2 flow channels in a polyether ether ketone flow cell, exposed to the metal side of the sensor. We used PBS as running buffer and ran at ∼5 – 20 µL/min during the experiments. Once a stable baseline was achieved, injections of necessary solutions were automated by using an autosampler and a 96-well plate. The temperature was kept at 25°C in all the experiments. The quantification of thickness, mass coverage or surface grafting density of layers, adhered on silica-coated gold sensors were estimated as reported before (see supplementary methods)[3–5]. In brief, the SPR spectra after silica coating were fitted with Fresnel models to retrieve the dry film thickness by using a custom MATLAB (MathWorks, Natick, MA, USA) code[6]. In liquid, the surface coverages (Γ) were estimated from independently determined parameters (e.g. molar refractivity dn/dC, plasmonic field decay length δ, etc.) as described previously[7], see next section.

The conditions for binding poly-His-tag spike onto NTA-bilayers in SPR experiments were as following: (i) the formation of SLB was done by the injection of SUVs containing 0.25% or 1% of NTA functionalized lipids at concentration of 100 µg/ml and a flow rate of 20 µL/min, (ii) spike was injected at a concentration of 18.33 µg/ml and a flow rate of 5 µl/min for 1 h.

**Quantification of spike optical mass by SPR**

For quantifying the number of absorbed biomolecules and thus surface grafting density from the recorded SPR sensogram signal ($\Delta\theta$) presented in Fig. S1B, we have used the following equation:

$$\Gamma = {}^{\Delta\theta\delta}/_{S_0 b}.$$

Here $b$ is the increase in refractive index per mass concentration of the binding species (0.18 ml/g for proteins and 0.17 ml/g for supported lipid bilayers). $S_0$ is the bulk sensitivity of the instrument (angular shift per refractive index increment), which was estimated from Fresnel models at 670 nm to be 131 degrees[3]. $\delta$ is the decay length of the evanescent field, which is calculated from the dispersion relation and the field distribution of the surface plasmons[8]. Considering experimentally determined parameters for the Cr and Au thickness and permittivity, we have $\delta$ = 228 nm for our system.

Using this model, from the SPR sensograms in Fig. S1B, we calculated the spike density on a bilayer with 0.25% molar concentration of NTA to be 62 ng/cm$^2$, which corresponds to a trimer area density of 920 trimers/μm$^2$. Assuming a spherical liposome of diameter 125 nm, this corresponds to ~45 trimer per particle, comparable to what was observed in SARS-CoV-2 virions[9,10].

**Quartz crystal microbalance with dissipation monitoring measurement**

Quartz crystal microbalance with dissipation monitoring (QCM-D) measurements were performed with an AWS X4 QCMD system (AWSensors, Valencia, Spain) using silica-coated sensors (AWS SNS 000049 A, AWSensors). The sensors were cleaned in SDS (2% w/v) for 30 minutes and then rinsed 2 times in 99.9% ethanol and 10 times in milli-Q water and then dried under nitrogen flow. The sample chamber and tubing were washed for 30 minutes in Cobas cleaner (20754765322, Roche, Basel, Switzerland), rinsed with abundant milli-Q water and dried under nitrogen flow. The sensor was then mounted into the chamber and kept under HBS flow. The supported lipid bilayer (SLB) was formed using POPC:DGS-NTA:LissRhod (99-X:X:1 molar ratio) vesicles at a final concentration of 100 μg/ml in 20 mM $NiCl_2$ in HBS. The vesicles spontaneously ruptured on the silica surface, forming an SLB. After rinsing the sample with HBS, 50 μl of spike solution (20-100 μg/ml) in HBS was injected; the protein solution was incubated for >30 min in static conditions. The chamber was then rinsed in PBS and the spike detached using a 0.5 M solution of imidazole in milli-Q water, to verify specific attachment via NTA-his-Tags. All solutions were injected at a flow rate of 20 μl/min. In between measurements, the sensors were cleaned in SDS and water, stored in SDS or dried over nitrogen flow, and reused up to 10 times.

The bound mass ($\Delta m$) including the water coupled with the protein layer was evaluated using the Sauerbrey model for a rigid film from the measured frequency shift ($\Delta f$) according to the equation: $\Delta m = -C\ \Delta f/n$. Where $C$ is the mass sensitivity constant and equals 17.7 ng/(cm$^2$·Hz) in our case, and $n$ is the number of the harmonic ($n = 3$). Due to the large water coupling and softness of the spike layer, the results of this model are only used as a coarse estimation. Refer to SPR measurements for a more accurate estimation of spike density[11].

**Cell binding assay**

Calu-3 cells were purchased from ATCC (HTB55, Manassas, VA, USA) and cultured in high-glucose DMEM (D5648, Sigma-Aldrich) 10-20% Fetal Bovine Serum (FBS, SV30160.03, HyClone, USA) and 1% Penicillin/Streptomycin (PenStrep, 10,000 Units/ml Penicillin, 10,000 μg/ml Streptomycin, 15140-122, Gibco). Four days prior to the experiment, cells were seeded on a glass-bottom 96-well plate at a density of ~10$^5$ cells/well. On the day of the experiment, cells were then washed 3 times in PBS and moved on ice, where they were incubated with 10 µl of spike-liposomes solution in 40 µl of DMEM + 3% FBS for 1 hour. They were then washed 3 times in ice-cold PBS and fixed in ice-cold 4% paraformaldehyde (PFA) for 10 min. The wells were washed 3 times in PBS and kept in PBS at 4°C until imaging. The wells were imaged on Nikon Ti2-E microscope in widefield configuration using a Spectra III solid-state light source (555 nm emission wavelength), a multiband pass filter cube 86012v2 DAPI/FITC/TxRed/Cy5 (Nikon Corporation, Melville, USA), a CFI Apochromat TIRF 60XC oil-immersion 60x objective (Nikon, NA: 1.49) and a Prime 95B sCMOS camera (Teledyne Photometrics, Birmingham, UK). A z-stack was acquired for each position, 5 per well, with step size 0.3 µm and range 25 µm. Brightfield images were also acquired. The fluorescent particles attached to cells were counted by performing a maximum intensity projection on the z-stack and then applying an in-house 2D peak-finder algorithm (see kinetic analysis section) to detect all the intensity peaks with prominence higher than 50.

**Production of native membrane vesicles from Calu-3 cells**

Calu-3 cells were grown in DMEM + 10% FBS + 1% PenStrep and expanded 1 in 4 roughly every week. 20 T-175 flasks were harvested to produce the stock used in this study using a protocol adapted from previous studies[12]. In brief, cells were washed 3 times in ice-cold PBS and then mechanically detached using a cell scraper in harvest buffer (PBS + protease inhibitor, cOmplete™, EDTA-free Protease Inhibitor Cocktail, 04693132001). The harvested cells were pelleted via centrifugation at 600×g for 10 min and then disrupted with a CF1 continuous cell disruptor (Constant Systems, UK). Nuclei and large organelles were pelleted at 2,000×g for 10 min, and the supernatant was collected and centrifuged at 6,000×g for 20 min to remove mitochondria. Finally, the native membrane vesicles (NMVs) were collected by centrifugation at 150,000×g for 90 min. Vesicle-containing plasma membrane material was purified using a sucrose gradient. The pellet was resuspended in harvest buffer and mixed with an equal volume of 80% weight/volume (w/v) sucrose solution. It was then layered with 30% w/v and 5% w/v sucrose solutions in harvest buffer and the gradient was centrifuged at 273,000×g for 2.5 h. NMVs raise to form a clear band between 5% and 30%, which was harvested, aliquoted, flash frozen, and stored at -80ºC. In all steps, the cellular material was maintained on ice or at 4ºC.

NMV material was quantified using a FRET assay previously developed by us and described previously[13].

**Formation of native supported lipid bilayer from Calu-3 cells**

Native supported lipid bilayers (nSLBs) from Calu-3 cells were formed as described previously[12]. In brief, NMVs from Calu-3 cells were mixed with PEG-POPC vesicles to a 12.5:87.5, diluted in PBS to a final concentration of 200 µg/ml and sonicated for 30 min at 40ºC in a bath sonicator (37 kHz, Elmasonic S40H, Germany). Tracer vesicles, 99:1 POPC:OG-DHPE, were then mixed to a 1:1000 weight ratio before nSLB formation. No.1 microscope borosilicate glass cover slides (diameter: 22 mm, VWR, 631-0158P) were cleaned by boiling for 2 h in a 10% v/v solution of 7x detergent (MP Biomedicals, CA) in milli-Q water, rinsed in abundant milli-Q water and stored in milli-Q water for up to 3 days. Before use, the slides were rinsed in milli-Q water, dried under nitrogen and treated in a UV/ozone oven (ProCleaner™ Plus, BioForce Nanosciences, Virginia Beach, VA, USA) for at least 30 min. Home-made PDMS sample holders with a sample volume of ~10 µl, cleaned in SDS, rinsed in milli-Q water and dried under nitrogen flow, were attached to the clean glass surface and 5 µl of PBS was added to each well. The vesicle mixture was added to the wells to a final concentration of 100 µg/ml. Bilayer formation was observed using TIRF microscopy or inspected after PBS rinsing for unruptured tracer vesicles.

The NMVs:PEG-POPC ratio was established by selecting the highest concentration of NMVs that ensure the formation of a continuous and mobile lipid bilayer with few to no tracer vesicles left after formation.

Enzymatic removal of HS from nSLB was performed by incubating the already formed bilayers with a cocktail of Heparinase I and III (2 units/ml each) in digestion buffer (20 mM Tris-HCl, 4 mM $CaCl_2$, 100 mM NaCl, 0.01% w/v BSA) for 1 hour at room temperature. The bilayers were then rinsed in PBS before the addition of spike-decorated liposomes.

**Production of nSLBs on glass beads and flow cytometry**

To produce nSLBs on silica beads, we adapted the protocol from the production of planar nSLBs. Acid-cleaned silica beads (microParticles GmbH, Berlin, Germany) were washed 3 times in PBS and then added to the vesicle solution, 5 µl of beads stock solution in 50 µl of hybrid vesicles, followed by incubation for 1 hour at room temperature under agitation. The beads were then washed 3 times in PBS and collected via centrifugation at 1,000×g for 1 minute. Enzymatic removal of HS was performed as described above, incubating the beads for 1 hour at 37°C. After washing in PBS, the beads were stained using anti-HS (370255-S, AMSBIO, United Kingdom) or anti-ΔHS antibody (370260-S, AMSBIO), which recognise heparan sulfate chains or the epitope resulting from heparinase cleavage respectively, at a concentration of 1:100 in 0.2% w/v BSA in PBS at room temperature for 2 h. After washing, the beads were incubated with a 1:200 dilution of alexa488-conjugated rabbit anti-mouse IgG/M (A-10680, Thermofisher Scientific) and Alexa488-conjugated anti-mouse IgG1 (A21202, Thermofisher Scientific) for the HS and ΔHS staining, respectively, in 0.2% w/v BSA in PBS for 2 hours. Finally, the beads were washed 3 times in PBS and the fluorescence signal was measured with a ZE5 cell Analyser (Bio-Rad). The flow cytometry data were analysed using FlowJo (BD, Franklin Lakes, NJ, USA).

**Immobilisation of glycosaminoglycans on supported lipid bilayers**

Borosilicate glass coverslips were prepared as for nSLB formation. SLBs were formed by spontaneous rupture of SUVs onto the clean glass surface. POPC:bioDOPE SUVs stock solution was diluted to a concentration of 100 µg/ml in PBS; 5 µl were added to each well and incubated for 30 min. The wells were rinsed, and streptavidin solution was added at a final concentration of 40 µg/ml. After 15 min of incubation, the wells were washed and incubated with 2% PFA for 10 minutes to crosslink streptavidin and prevent lateral diffusion of streptavidin across the bilayer. After rinsing, end-on biotinylated GAGs were added at a concentration of 40 µg/ml and incubated for 30 min to achieve full coverage of the surface. Finally, the wells were rinsed before the addition of spike-liposomes.

All steps were performed at room temperature and all washing steps consisted of removing 5 µl of solution and washing 7 times with 10 µl of PBS, always leaving 5 µl of solution in the well to prevent it from drying.

**Imaging and analysis of particle's surface attachment**

Particle attachment and detachment to SLBs were imaged by acquiring time-lapses with a Nikon Ti2-E microscope in TIRF mode (see Cell binding assay). After the bilayer formation and functionalisation, 5 µl of fluorescent spike-decorated liposomes (1% DGS-NTA) were added to 5 µl of PBS in each well and incubated for at least 1 hour to let the system reach equilibrium. Images (704 x 704 pixels, with a resolution of 0.183 µm/pixel) were acquired for 1.5 to 3 hours, in 4 randomly selected positions per well. The frame rate was set for each experiment to 25-40 seconds per frame. All wells were imaged simultaneously.

Image registration was performed with an in-house MATLAB script to remove subtle movements due to the parallel acquisition of several positions[14,15]. Particles were then tracked in the stabilized movies using an additional in-house MATLAB script as described previously[14]. In short, after a Gaussian smoothing ($\sigma = 1$ pixel) to reduce noise, a 2D peak finding algorithm was applied to each frame and only peaks with a prominence exceeding a threshold intensity of 50 were considered. The threshold was set by the user after visual inspection of the particle detection results and used for all the experiments. The detected peaks in successive frames are matched using a global nearest neighbour algorithm to determine the arrival and residence time of each particle. After tracking, duplicate traces are automatically removed and fragmented traces linked, and finally false detachment events due to fluorophore bleaching, i.e. a slow reduction of the particle intensity below the detection level, are removed from the analysis.

The remaining traces were analysed for the determination of the kinetic parameters of the bond, the particle binding rate and the association, $k_{on}$, and dissociation rate constant, $k_{off,}$ using equilibrium fluctuation analysis[15,16]. The relative variation of $k_{on}$ was determined by fitting the cumulative arrival time of the particle on the surface to a straight line: $y = r_a t + y_0$, where $r_a$ is the particle arrival rate and $y_0$ considers the particles already bound at the beginning of the acquisition ($t = 0$). Once normalised by the particle concentration in solution, the measured arrival rate, $k_m$, is proportional to $k_{on}$ and it can be used to compare the attachment rates between samples. $k_{off}$ was calculated by fitting the number of attached particles as a function of the residence time on the surface to a single exponential function and an offset ($C$), $y = A \exp(-k_{off} t) + C$. While the interaction of the particle with the surface might be heterogeneous, with the creation of multiple bonds and bonds to different species present on the bilayer, the distribution of the survival times of the particles on the surface can be approximated to a single exponential decay. This should be interpreted as an average of the possible interactions experienced by the particle. Only the particles landing in the first half of the movie were considered to avoid bias towards short residence times. Particles that did not detach after more than 2500 s were considered to be irreversibly bound to the substrate in the time scale of the experiment. The percentage of irreversibly bound particles over the total number of detected particles was defined as the irreversible fraction.

The particle concentration in solution was measured for every sample using "bouncing particle analysis" as described previously[14]. In brief, the sample was first bleached to remove the signal from particles on the surface, and then a 30-60 s video was acquired at 10 fps in TIRFM configuration. The videos were analysed in the same way as for the kinetic analysis, but, in this case, only particles entering the TIRFM volume but not binding to the surface (residence time lower than 1 s) were considered. Since the particles freely diffuse in the buffer, the number of the recorded particles is proportional to their concentration in solution. In this way, the relative concentration of the particles between samples can be measured. The linearity between "bouncing particle analysis" results and particle concentration was demonstrated by measuring a serial dilution of fluorescently labelled SUVs on an inert POPC bilayer, Fig. S11. The SUV stock was diluted in PBS 100, 200, 400 and 800 times and three areas were imaged per condition.

All kinetic parameters are reported as normalised to the value obtained for Omicron for each sample. In this way, we accounted for the experimental variations due to changes in the liposome and bilayer composition between experiments. A summary of the non-normalised data can be found in Table S1-3.

**Preparation of HS sample for atomic force microscopy-based force spectroscopy**

Microscopy glass cover slides with a diameter of 24 mm (VMR, No. 2) were cleaned with 7x detergent and milli-Q water with 1:6 v/v solution at 70°C for 2 hours, rinsed thoroughly, and stored in milli-Q water. The slides were rinsed with excessive milli-Q water, nitrogen dried, and treated in UV/ozone cleaner for 30 min before use.

A cleaned glass coverslip was fixed on a stainless-steel atomic-force microscopy (AFM) holder (JPK/Bruker) using a bi-component Twinsil glue (Picodent, Germany) creating a well that holds a maximum volume of about 200 µL for preparing SLBs. SLBs were formed by spreading SUVs (100 µL of 100 µg/ml of POPC:bioDOPE; 90:10 molar ratio mixture) in the well and incubating for 30 mins. After forming SLBs, the well was rinsed by adding and removing 100 µl of PBS 20 times. Next, streptavidin was added to the well at the final concentration of 50 µg/ml (total volume 200 µl) and incubated for 20 mins. After the rinsing step (30 times with PBS), the streptavidin-coated SLB was exposed to biotinylated HS at the final concentration of 4 µg/ml for 30 mins. For biotinylated HA, the final concentration was increased to 7 µg/ml while keeping the same incubation time. All incubation steps were performed under stagnant conditions. The solution was homogenized at each dilution step with at least 2 times aspiration and release of ~50 µl of solution using a pipette.

**Functionalization of AFM tip with spike or its variants**

The AFM tip was functionalised with a PEG linker following the protocol in [17,18]. Briefly, AFM probes (MSCT, Bruker) were treated in UV/ozone cleaner for 30 mins. The cleaned cantilevers were then exposed to APTES (3-Aminoproyl)triethoxysilane) and triethylamine using the gas-phase silanization reaction[19]. Next, the cantilevers were incubated in 0.5 ml of chloroform containing 1 mg of NHS-PEG-Acetal MW 2k (Creative PEG, USA) and 30 µL of triethylamine for 2 hours. Following washing with chloroform (5 min, 3x), the cantilevers were dried with nitrogen and either stored under nitrogen until use or functionalized instantly with Tris-NTA-amine linker (BR1001101, biotechrabbit, Germany). For the functionalization with Tris-NTA-amine linker, one of the cantilevers was immersed in 1% w/v citric acid in milli-Q water for 10 mins. After washing with milli-Q water (5 min, 3x), the cantilever was incubated in the solution containing 0.71 µg/ml of Tris-NTA-amine in PBS and 2 µL of freshly prepared sodium cyanoborohydride ($NaCNBH_3$) solution (13 mg of $NaCNBH_3$ dissolved in 20 µL of 20 mM NaOH and 180 µL of milli-Q water) for 30 mins. The coupling reaction was then quenched with 5 µL of 1 M (pH 8.0) ethanolamine solution and incubated for 10 min. The cantilever was then carefully rinsed 4 times with 1 ml PBS, dried with nitrogen, and instantly mounted on the AFM cantilever holder. Next, 100 µL of 10 mM $NiCl_2$ in HEPES buffer was carefully added to the cantilever and incubated for 15 mins. After rinsing twice with 1 ml HEPES and twice with 1 ml PBS, the cantilever was then incubated with His-tagged spike at the final concentration of 1.5 µg/ml for 10 mins to ensure the coupling of a very low concentration of spike proteins on the tip apex. All chemicals used for AFM tip functionalization were purchased from Sigma-Aldrich unless otherwise stated.

**AFM-SMFS experiments**

All SMFS experiments were performed in PBS at room temperature using a JPK/Bruker 4XP BioScience with the JPK SPM Control Software v.7 (JPK/Bruker, Berlin, Germany). Force-distance (FD) curves were recorded using the cantilevers with nominal spring constants of 10 pN/nm (tip C) and

30 pN/nm (tip D) of the MSCT probes. Before recording FD curves, the real spring constants of each tip were measured using the thermal noise method[20] and they were determined to be within 10% of the nominal values given by the supplier. To determine the dynamics of the spike-HS binding interactions, the spike-functionalized tip was moved toward the HS surface (ramp size = 400 nm) at the speed of 1 µm/s until in contact with the surface (force load threshold of 600 pN). The tip was left interacting with the surface (dwell time) and then retracted. For dissociation rate constant ($k_{\mathrm{off}}^{\mathrm{sm}}$) measurements, retract speeds were varied from 1 to 7 µm/s, and 1500 to 2500 FD curves were recorded by keeping the approach speed fixed to 1 µm/s and no dwell time. For association rate constant ($k_{\mathrm{on}}^{\mathrm{sm}}$) measurements, both approach and retract speeds were set to 1 µm/s to probe the probability of bond formation (binding probability, BP) as a function of dwell time. For this, 400-500 FD curves were recorded for dwell times of 50, 100, 150, 200, 300, 500, and 1000 ms. For all spike variants, at least 2 experiments with independently prepared tips and surfaces were performed unless stated otherwise. To ensure the single-molecule nature of the interaction, conditions for functionalization of both the AFM tip and the HS surface were adjusted such that the binding probability of unbinding events was <20%.

**SMFS data analysis**

All SMFS data were processed using the JPK data processing software (JPK/Bruker, Germany), OriginPro 2016 (OriginLab Corporation, USA), and GraphPad (Prism 9, USA). The retract curves were analysed using the freely joint chain (FJC) model, which approximates the polymer to a chain of freely joined rigid segments. To identify the stretching of the PEG linker and rupture (unbinding) event for spike-HS bond, each curve was fit keeping both contour (total or end-to-end length of a chain) and Kuhn (a measure of elasticity of a polymer chain, equal to the length of each segment) lengths as fitting parameters[21]. The curves with unbinding events occurring at less than 10 nm from the surface were discarded to avoid the contribution from non-specific interactions. The binding probability was calculated as the fraction of curves displaying rupture events. From the FJC fitting, rupture force and instantaneous loading rate (*r*) were determined by multiplying the effective spring constant, i.e., the slope of the FJC fit close to the rupture point, by the retract speed. Force histograms were constructed to calculate the mean rupture force from every unbinding event at the same retract speed or loading rate and fitted with a Gaussian function in Origin (OriginLab Corp., Northampton, MA, USA). Next, the position of the fit maxima, which reflects the most probable rupture force (*F*), was plotted as a function of the instantaneous loading rate (*r*) to create dynamic force spectra (DFS) plots. DFS data was then fitted with the Bell-Evan model ($F = \frac{k_{\mathrm{B}}T}{x_{\beta}} \ln\left(\frac{\mathrm{r}x_{\beta}}{k_{\mathrm{off}}^{\mathrm{sm}} k_{\mathrm{B}}T}\right)$) to extract the intrinsic bond kinetic parameters (i.e. $k_{\mathrm{off}}^{\mathrm{sm}}$ and $x_{\beta}$) using linear regression analysis, where $x_{\beta}$ is the width of the energy barrier and $k_{\mathrm{off}}^{\mathrm{sm}}$ is the rate constant at $F = 0$, $k_{\mathrm{B}}T$ is the product of Boltzmann constant and the temperature[22–24]. Error bars in DFS plots represent the standard deviation of the Gaussian fit of mean rupture force histograms.

To determine the single bond association rate, $k_{\mathrm{on}}^{\mathrm{sm}}$, the binding probability was plotted as a function of dwell time, $t$, (i.e., the time tip stays in contact with the surface or the time between the end of the approach and the beginning of the retraction steps) and fitted with the following pseudo-first-order kinetics function: $\mathrm{BP} = \mathrm{A}\left[1 - \exp\left(-\frac{t-t_0}{\tau}\right)\right]$, where A is the maximum measured BP, $t_0$ is the lag time, and τ is the interaction time between spike and HS. $k_{\mathrm{on}}^{\mathrm{sm}}$ was determined from following function: $k_{\mathrm{on}}^{\mathrm{sm}} = \tau^{-1}{\mathrm{C}_{\mathrm{eff}}}^{-1}$, where $\mathrm{C}_{\mathrm{eff}}$ is effective concentration. The effective concentration was calculated using the following expression[25,26]; $\mathrm{C}_{\mathrm{eff}} = 3n_b/2\pi {r_{\mathrm{eff}}}^3 \mathrm{N}_\mathrm{A}$, where $r_{\mathrm{eff}}$ is the radius of the sphere with volume $\mathrm{V}_{\mathrm{eff}}$ and it is assumed to be equal to the equilibrium length of the PEG linker, ~2 nm, plus the size of spike, ~20 nm, $n_\mathrm{b}$ is the number of potential bonds formed within the hemisphere in which interaction can take place (this is set to 1 in our case, as majority of FD curve show only a single unbinding event) and $\mathrm{N}_\mathrm{A}$ is the Avogadro's number. From the $k_{\mathrm{off}}^{\mathrm{sm}}$ and $k_{\mathrm{on}}^{\mathrm{sm}}$ values, the binding affinity ($\mathrm{K}_\mathrm{D}^{\mathrm{sm}}$) was calculated and mean ± standard deviation was reported. The standard deviation was determined by propagating the experimental error through the abovementioned formulas.

**Safety Statement**

No unexpected or unusually high safety hazards were encountered.

### Supplementary figures

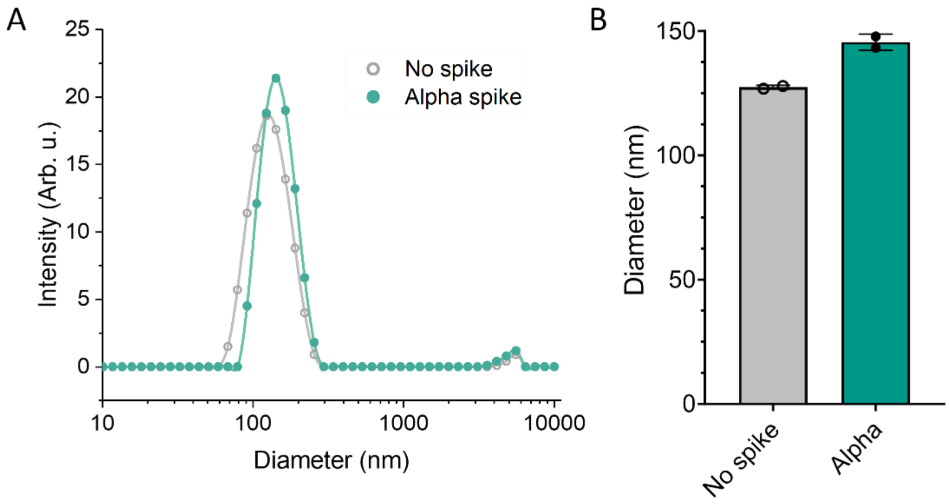


**Figure S1 A.** Size distribution of naked NTA-liposomes and liposomes decorated with soluble spike (Alpha variant) measured by dynamic light scattering (DLS). Solid lines are the cubic spline interpolation of the experimental data (dots) **B.** Mean diameter of naked and spike-decorated liposomes calculated from DLS measurements.

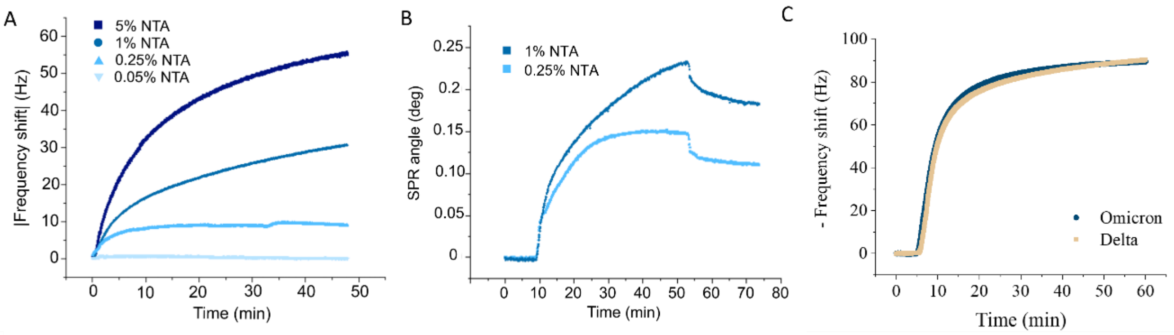


**Figure** S2 **A**. Absolute value of the QCM-D frequency shift due to the binding of soluble His-tagged spike on an NTA-presenting SLB, showing the dependency of spike binding to the concentration of NTA-conjugated lipids. Molar fraction of NTA-lipids between 0.05% to 5%. **B.** SPR measurement of spike attachment to NTA-presenting SLBs with a molar fraction of NTA-lipids of 0.25% and 1% used to estimate the spike density on the particles. **C.** Frequency shift of the attachment of spike from Omicron and Delta variants onto an NTA-bilayer (molar fraction: 1%). The nearly identical profile indicates that the protein loading is independent of the SARS-CoV-2 variant.

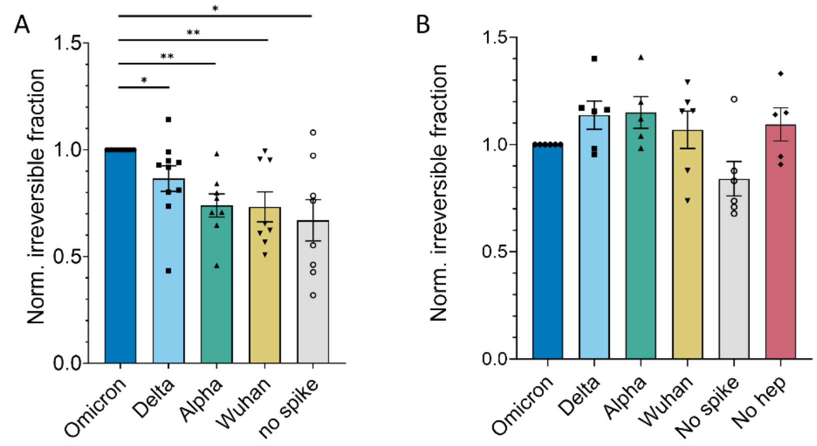


**Figure S3** Fraction of spike-decorated liposomes forming a bond with a lifetime longer than 2500 s on untreated Calu-3-derived nSLBs (**A**) and nSLBs treated with heparinase I and III (**B**). Data are normalised by the value measured for Omicron in each experiment. Statistical significance calculated using one-way ANOVA test. *: $p<0.05$, **: $p<0.01$.

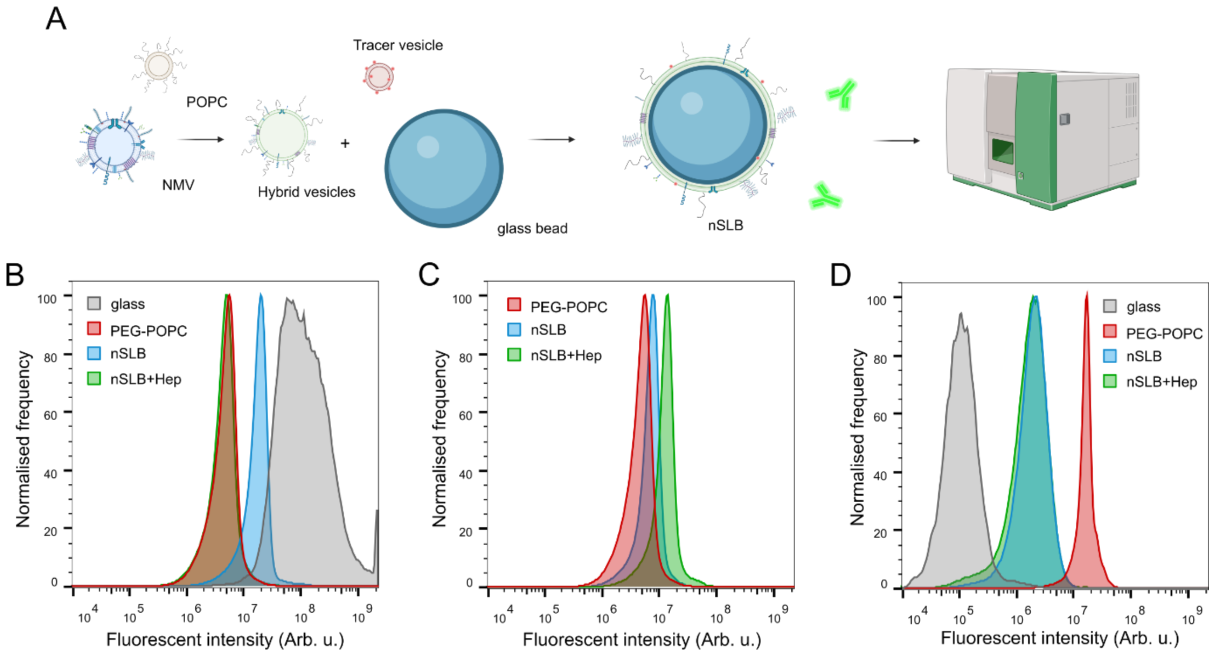


**Figure S4 A.** Schematics showing the process of forming nSLB on glass beads, antibody staining and final readout using flow cytometry. **B.** Flow cytometry signal from antibody staining of nSLB-coated beads for heparan sulfate, showing the signal reduction to the level of the negative control (PEG-POPC) after heparinase treatment (nSLB+Hep). Naked glass beads show a high signal due to non-specific absorption of the secondary antibody. **C.** Flow cytometry signal from antibody staining of nSLB-coated beads against the cleavage site of heparinase-treated heparan sulfate. **D.** Flow cytometry signal from the fluorescent lipids incorporated into the nSLB, showing the presence of the nSLB on beads. Fluorescent lipids are incorporated by adding vesicles containing 1% OG-DHPE to the PEG-POPC:NMV solution in a 1:500 v/v ratio. Pure PEG-POPC bilayers without the addition of membrane material appear brighter due to more efficient incorporation of tracer vesicles during the bilayer formation.

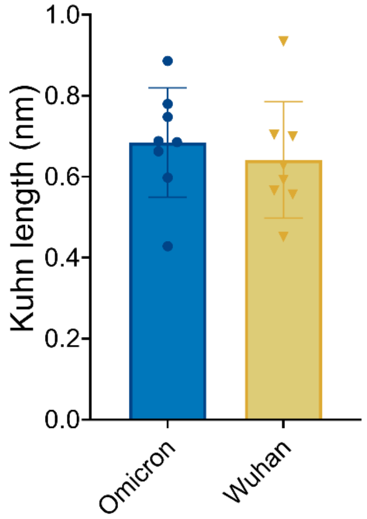


**Figure S5** Kuhn lengths of the PEG chain used to immobilise spike variants on the AFM tips for SMFS experiments. Average values of 0.68 ± 0.13 (Omicron) and 0.64 ± 0.14 nm (Wuhan).

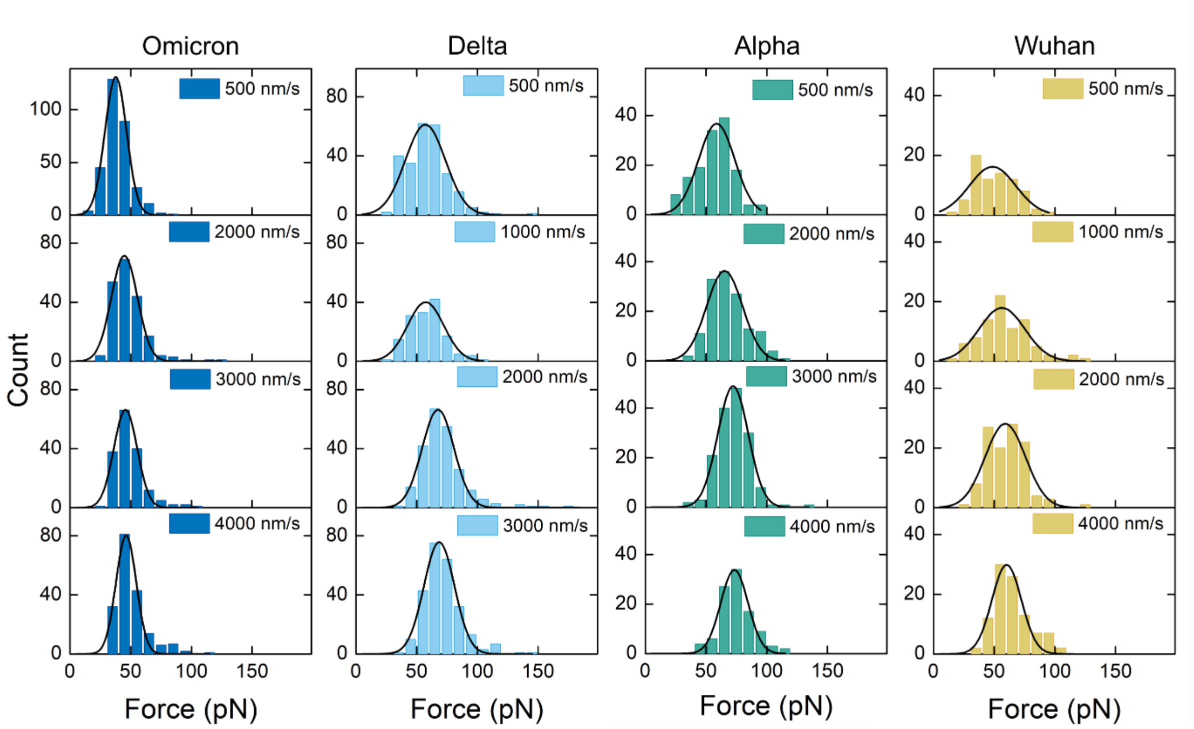


**Figure S6** Rupture force distribution for spike-HS interactions for four variants as obtained by AFM-based SMFS.

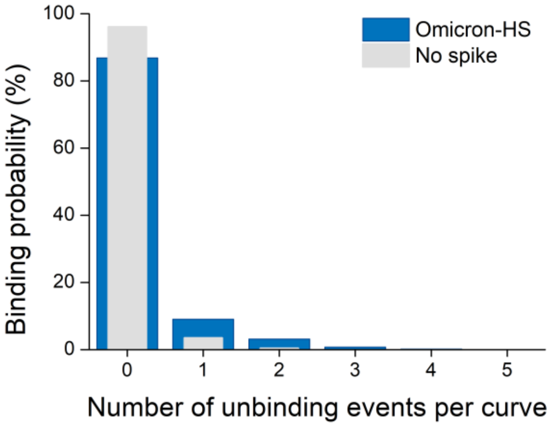


**Figure S7** Binding probability for Omicron-HS (blue) and NTA-HS (light grey) in SMFS experiments.

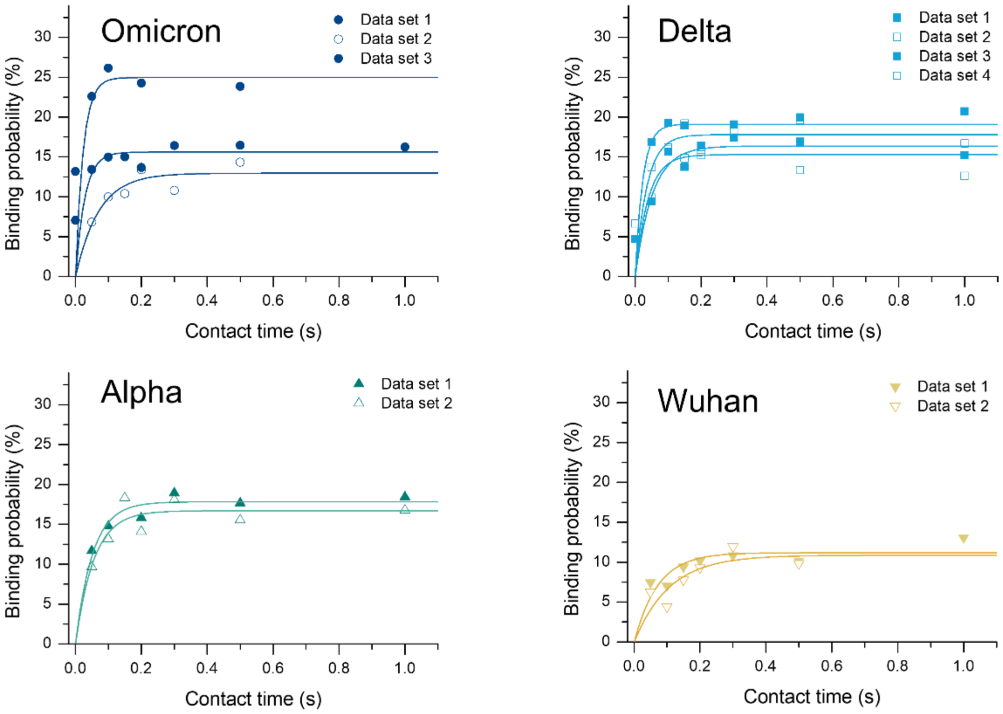


**Figure S8** Binding probability data as a function of contact time for all four variants used to determine interaction time (τ) by fitting binding probability with a mono-exponential function (solid lines).

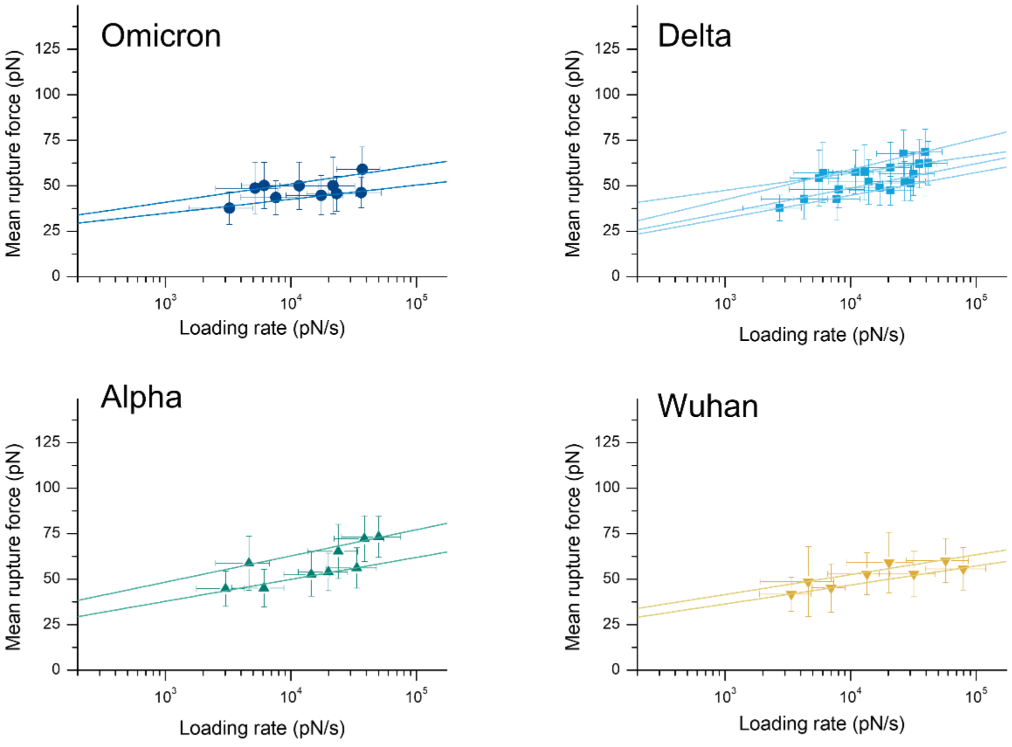


**Figure S9** DFS data sets for all four variants with error bars (sigma from Gaussian fit of force histograms) to determine $k_{\mathrm{off}}$ and $x_\beta$ by fitting DFS according to the Bell-Evans model (solid lines).

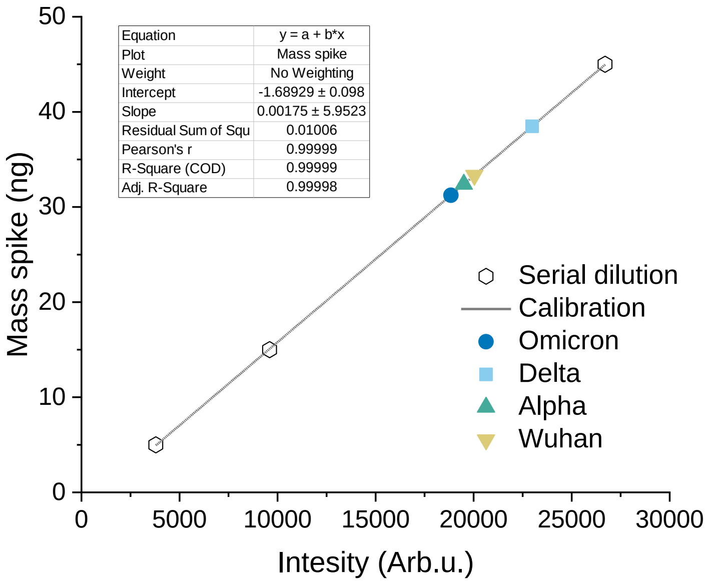


**Figure S10** Calibration curve used to measure the spike content in spike-decorated liposomes lysate from the western blot in Fig. 1D.

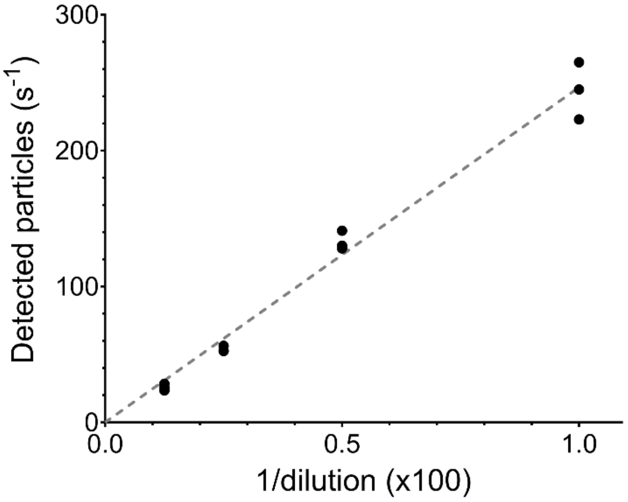


**Figure S11** Number of newly detected particles per second in a serial dilution of fluorescently labelled liposomes on a POPC bilayer. The black dots show the experimental data and the dashed grey line the linear fit passing through the origin ($y = 246.6x$, R squared: 0.9821).

#### Supplementary tables

| | Omicron | Delta | Alpha | Wuhan | No spike |
|---|---|---|---|---|---|
| $k_{\mathrm{m}}$ (Arb.u.) | 1.61 (0.34) | 0.94 (0.11) | 1.30 (0.49) | 1.51 (0.50) | 0.28 (0.08) |
| $k_{\mathrm{off}}$ ($s^{-1}$) ($x10^{-4}$) | 4.14 (0.62) | 8.45 (2.33) | 4.75 (0.78) | 4.49 (0.71) | 27.0 (8.43) |
| Irreversible fraction (%) | 53.8 (4.3) | 42.9 (4.7) | 47.7 (4.2) | 48.1 (4.7) | 33.0 (4.4) |

**Table S1** Mean and standard error of the mean, in brackets, of the kinetic parameters of the multivalent interaction between spike-decorated liposomes and Calu-3 derived nSLBs, prior to the normalisation to the Omicron sample used in Fig.3 and Fig. S3A.

| | Omicron | Delta | Alpha | Wuhan | No spike | Omicron (no hep.se) |
|---|---|---|---|---|---|---|
| $k_{\mathrm{m}}$ (Arb.u.) | 2.78 (0.44) | 1.98 (0.32) | 7.79 (2.06) | 4.61 (0.14) | 0.67 (0.14) | 2.59 (0.55) |
| $k_{\mathrm{off}}$ ($s^{-1}$) ($x10^{-4}$) | 8.30 (1.78) | 13.3 (6.12) | 8.49 (2.53) | 10.4 (4.43) | 25.4 (3.87) | 5.44 (0.68) |
| Irr. fraction (%) | 50.1 (2.9) | 54.3 (1.7) | 55.3 (3.3) | 51.3 (3.4) | 41.3 (4.4) | 52.6 (3.2) |

**Table S2** Mean and standard error of the mean, in brackets, of the kinetic parameters of the multivalent interaction between spike-decorated liposomes and heparinase treated Calu-3 derived nSLBs, prior to the normalisation to the Omicron sample used in Fig.4 and Fig. S3B.

| | Omicron | Delta | Alpha | Wuhan | HA |
|---|---|---|---|---|---|
| $k_m$ (Arb.u.) | 4.92 (1.30) | 0.50 (0.18) | 0.27 (0.05) | 0.19 (0.05) | 0.08 (0.04) |
| Surface coverage (part./frame) | 2987 (777) | 283 (73) | 100 (19) | 95 (22) | 33 (13) |

**Table S3** Mean and standard error of the mean, in brackets, of the kinetic parameters of the multivalent interaction between spike-decorated liposomes and SLB-immobilised heparan sulfate prior to the normalisation to Omicron used in Fig.5
